# Divergence, gene flow and the origin of leapfrog geographic distributions: The history of color pattern variation in *Phyllobates* poison-dart frogs

**DOI:** 10.1101/2020.02.21.960005

**Authors:** Roberto Márquez, Tyler P. Linderoth, Daniel Mejía-Vargas, Rasmus Nielsen, Adolfo Amézquita, Marcus R. Kronforst

## Abstract

The geographic distribution of phenotypic variation among closely related populations is a valuable source of information about the evolutionary processes that generate and maintain biodiversity. Leapfrog distributions, in which phenotypically similar populations are disjunctly distributed and separated by one or more phenotypically distinct populations, represent geographic replicates for the existence of a phenotype, and are therefore especially informative. These geographic patterns have mostly been studied from phylogenetic perspectives to understand how common ancestry and divergent evolution drive their formation. Other processes, such as gene flow between populations, have not received as much attention. Here we investigate the roles of divergence and gene flow between populations in the origin and maintenance of a leapfrog distribution in *Phyllobates* poison frogs. We found evidence for high levels of gene flow between neighboring populations but not over long distances, indicating that gene flow between populations exhibiting the central phenotype may have a homogenizing effect that maintains their similarity, and that introgression between “leapfroging” taxa has not played a prominent role as a driver of phenotypic diversity in *Phyllobates*. Although phylogenetic analyses suggest that the leapfrog distribution was formed through independent evolution of the peripheral (i.e. leapfrogging) populations, the elevated levels of gene flow between geographically close populations poise alternative scenarios, such as the history of phenotypic change becoming decoupled from genome-averaged patterns of divergence, which we cannot rule out. These results highlight the importance of incorporating gene flow between populations into the study of geographic variation in phenotypes, both as a driver of phenotypic diversity and as a confounding factor of phylogeographic inferences.

## Introduction

Geography has a strong influence on the diversification of closely related lineages, since it largely mediates the level of gene flow between them (Huxley, 1942; Mayr, 1942). Therefore, studying the geographic distribution of phenotypic and genetic variation among such lineages can generate valuable insights into the processes that generate biological diversity. An intriguing pattern of geographic variation is the “leapfrog” distribution, where phenotypically similar, closely related populations (of the same or recently diverged species) are disjunctly distributed and separated by phenotypically different populations to which they are also closely related (Chapman, 1923; Remsen, 1984). Such patterns have been reported in multiple taxa, such as birds (e.g. Cadena, Cheviron, & Funk, 2010; Chapman, 1923; Norman, Christidis, Joseph, Slikas, & Alpers, 2002; Remsen, 1984), flowering plants (Matsumura, Yokohama, Fukuda, & Maki, 2009; Matsumura, Yokoyama, Tateishi, & Maki, 2006), and butterflies (Brower, 1996; Emsley, 1965; Hovanitz, 1940; Sheppard, Turner, Brown, Benson, & Singer, 1985). Since leapfrog patterns represent repeated instances of similar phenotypes in space, they provide a rich opportunity to understand the processes generating phenotypic geographic variation.

Two main hypotheses have been put forward to explain the origin of leapfrog distributions (Norman et al., 2002; Remsen, 1984): First, the phenotypically similar, geographically disjunct populations can owe their resemblance to recent common ancestry (i.e. they are descendants of an ancestral population with the same phenotype), and the disjunct range of “leapfrogging” forms is due to biogeographic processes such as long-range migration or the extinction of geographically intermediate populations. Second, the distribution of phenotypes may be due to evolutionary convergence of populations with similar phenotypes, or divergence of the central (intervening) populations from the ancestral phenotype. Clear phylogenetic predictions can be drawn from these hypotheses: If phenotypic similarity among the leapfrogging populations is due solely to recent common ancestry, then such populations should be more closely related to one another than to geographically close populations that display the intervening phenotype. If the geographic distribution of phenotypes is due to convergent or divergent evolution then a correspondence between phylogeny and phenotypes is not expected. In this case, however, ancestral state reconstructions can identify whether the central or peripheral populations exhibit derived (i.e. divergent) phenotypes. Accordingly, efforts to elucidate the evolutionary mechanisms behind leapfrog distributions have mainly focused on inferring the phylogenetic relationships among populations and using them to reconstruct the evolution of the phenotype in question (e.g. Brower, 1996; Cadena et al., 2010; Shun’Ichi Matsumura et al., 2009; Norman et al., 2002; Quek et al., 2010).

Although a cladogenetic description of population history can reveal a great deal about the origin of leapfrog distributions, it is unable to capture some important aspects of the diversification process. Among them is the extent of gene flow between populations (or its absence), which can play an important role in the formation of leapfrog distributions. For instance, reduced levels of genetic exchange between populations with different phenotypes will promote the existence of such differences, while introgressive hybridization between populations can facilitate phenotypic similarity between them. Furthermore, if gene flow between geographically close populations with different phenotypes is pervasive, it can homogenize previous genetic divergence between these populations, decoupling the history of the phenotype from genome-wide patterns of divergence (Hines et al., 2011; James, Arenas-Castro, Groh, Engelstaedter, & Ortiz-Barrientos, 2020), which can complicate inferences related to the origin of leapfrog distributions.

Here we examine the processes driving the origin of a leapfrog distribution present in *Phyllobates* poison-dart frogs. This genus is found from Southern Nicaragua to Western Colombia, and is composed of five nominal species (Myers, Daly, & Malkin, 1978; Silverstone, 1976): *P. vittatus*, *P. lugubris*, and *P. aurotaenia,* which exhibit a bright dorsolateral stripe on a dark background, and *P. terribilis* and *P. bicolor*, which display solid bright-yellow dorsal coloration (Fig. 1A). The two latter species exhibit a leapfrog distribution in Western Colombia, separated by *P. aurotaenia*: *P. bicolor* occurs on the slopes of the Western Andes, in the upper San Juan river basin, *P. aurotaenia* in the lowlands along the San Juan and Atrato Drainages and onto the Pacific coast, and *P. terribilis* along the Pacific coast south of the San Juan’s mouth (Fig. 1C).

**Fig 1.**
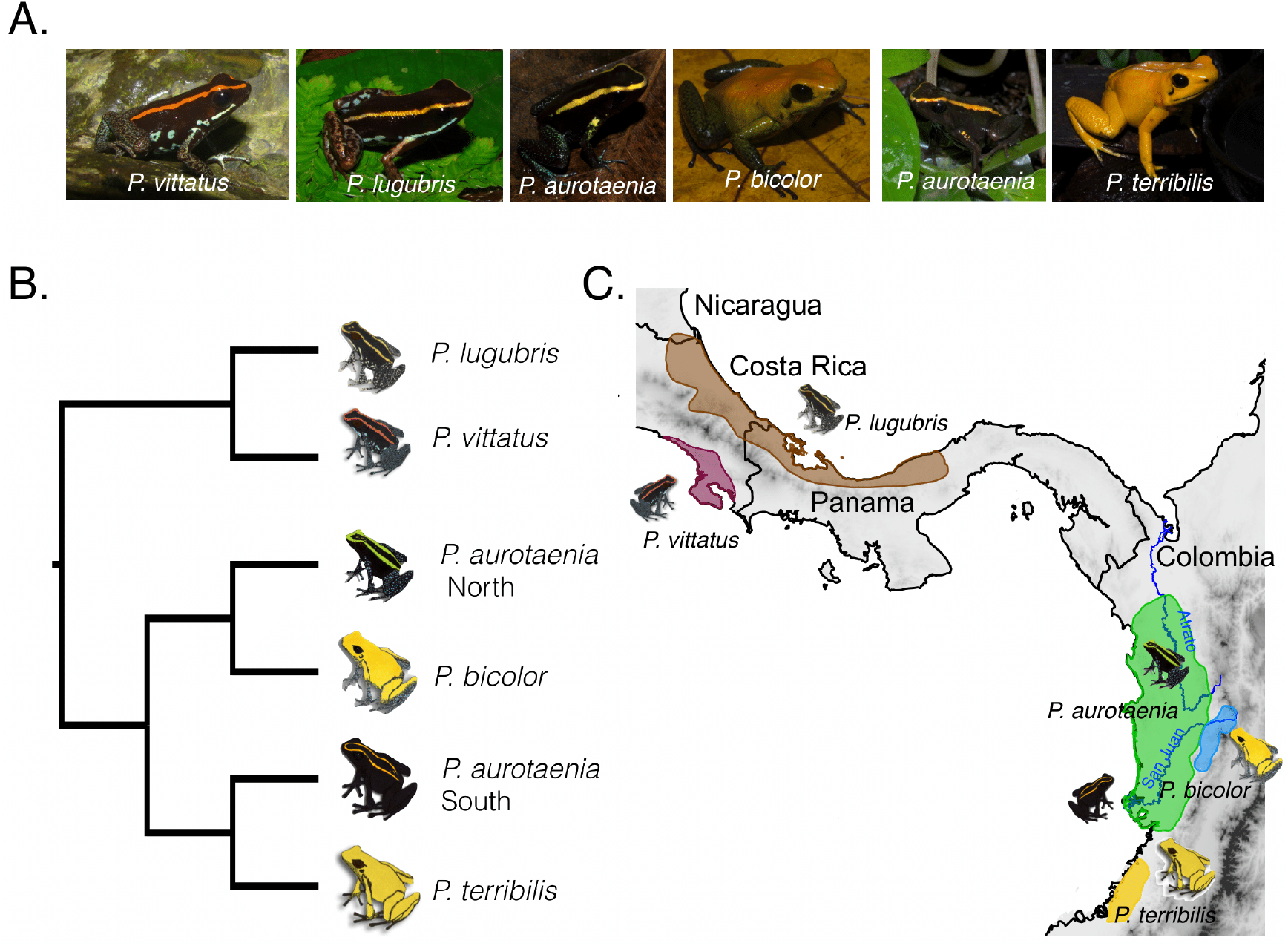
A) Color pattern diversity, B) currently accepted phylogenetic relationships (Grant et al., 2017), and C) geographic distribution of *Phyllobates* poison frogs. Species distribution polygons were obtained from the IUCN red list of threatened species website (https://www.iucnredlist.org/) and modified to fit natural history collection records and our own observations.

Early systematic work grouped *P. terribilis* and *bicolor* as sister species based on morphological and ontogenetic characters (Maxson & Myers, 1985; Myers et al., 1978). Although an early mitochondrial phylogeny supported these relationships (Widmer, Lötters, & Jungfer, 2000), subsequent work has consistently recovered *P. terribilis* and *P. bicolor* as non-sister taxa (Grant et al., 2006, 2017; Márquez, Corredor, Galvis, Góez, & Amézquita, 2012; Santos et al., 2009), and even suggested that *P. aurotaenia* may actually represent two separate lineages, one sister to *P. bicolor* and the other to *P. terribilis* (Grant et al., 2017; Santos et al., 2009). Although these studies only included 1-4 samples per *Phyllobates* species, and were based on DNA sequences from a small number of markers (1-7 loci), their results are compatible with convergent evolution giving rise to the leapfrog distribution.

In this study we aim to shed light on the evolutionary genetic and biogeographic processes involved in the origin of the current geographic distribution of aposematic coloration in *Phyllobates* poison frogs. Based on substantially increased sampling across Colombian populations and thousands of genome-wide markers, we leverage phylogenetics and spatial population genetics to 1) elucidate the extent of genetic structure and evolutionary relationships among populations, and 2) evaluate the role of gene flow between populations in the formation of the leapfrog distribution.

## Materials and Methods

To obtain a representative sample of Colombian *Phyllobates* populations, we conducted field expeditions to 23 localities throughout the genus’s range (Fig. 2B), resulting in tissue (i.e. mouth swab, toe-clip, or liver) samples from 108 individuals (Table S1). In addition, we obtained eight samples of *P. vittatus* and *P. lugubris*, (four samples per species; Table S1) to serve as outgroups in our analyses. Both species are distributed in Central America, and have been consistently found to be the sister group of Colombian *Phyllobates* (Grant et al., 2006, 2017; Santos et al., 2009; Widmer et al., 2000).

**Fig 2.**
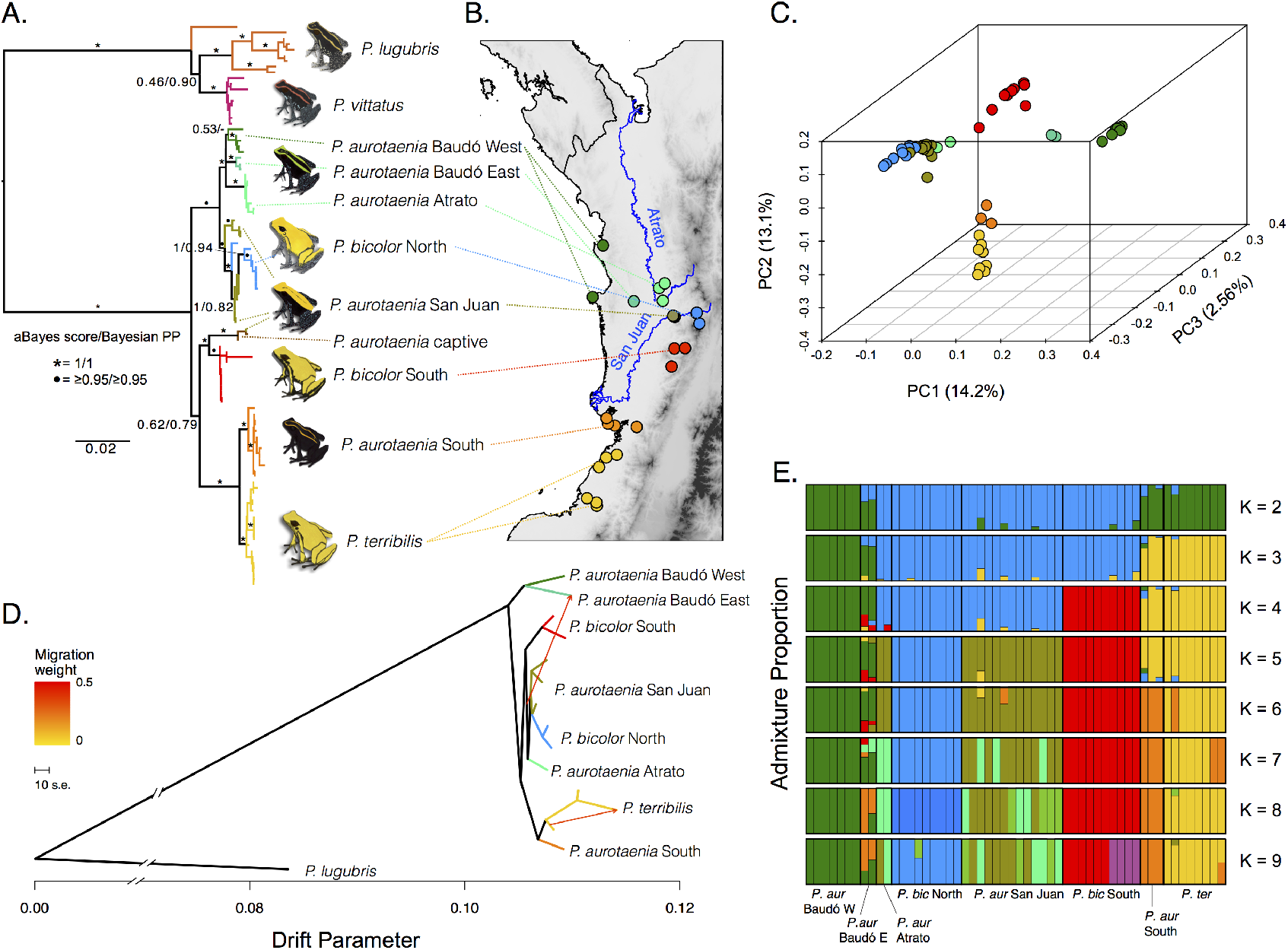
Genetic structure among *Phyllobates* populations in western Colombia. A) Maximum likelihood mtDNA genealogy inferred from 1926bp. B) Sampling localities for this study. C) Principal component analysis plot based on the first three components accounting for 29.4% of the variance. D) Treemix population graph assuming 2 migration edges. E) Individual admixture proportions assuming 2-9 ancestral populations. Colors in A-D correspond to the operational taxonomic units (OTU) used for phylogenetic analyses. Colors in E were chosen to loosely represent these clusters.

### mtDNA Sequencing and analysis

To gain initial insight into the levels of genetic variation and structure among populations we sequenced fragments of three mtDNA markers: 16S rRNA (16S; 569bp), Cytochrome Oxidase I (COI barcoding fragment; 658bp), and Cytochrome b (Cytb; 699bp) from 74 individuals. We extracted DNA using either Qiagen DNeasy spin columns or a salt precipitation protocol (Miller, Dykes, & Polesky, 1988), and used primers 16Sar and 16Sbr (Palumbi et al., 1991), Chmf4 and Chmr4 (Che et al., 2012), and CytbDen3-L and CytbDen1-H (Santos & Cannatella, 2011) to amplify the 16S, COI, and Cytb loci, respectively. Thermal cycling protocols consisted of 2 min at 95°C, 30-35 cycles of 30 sec at 95°C, 1 min at 45°C and 1.5 min at 72°C, and a final 5 min at 72°C. PCR products were purified with ExoSAP (Affymetrix) and sequenced in both directions using an ABI 3500 Genetic Analyzer (Applied Biosystems). Chromatograms were assembled and visually inspected in Geneious R9 (Kearse et al., 2012) to produce finalized consensus sequences.

We aligned our sequences and those available in GenBank (Table S1) using MUSCLE (Edgar, 2004), and built mtDNA trees with PhyML 3.3 (Guindon et al., 2010) and MrBayes 3.2.6 (Altekar, Dwarkadas, Huelsenbeck, & Ronquist, 2004; Ronquist et al., 2012). MrBayes analyses consisted of 10 million iterations (two runs with four chains each), sampling every 1,000 iterations, and discarding the first 2,500 trees (25%) as burnin. PhyML runs started from five different random trees, and used SPR moves to search the tree space. Nodal support was evaluated using aBayes scores (Anisimova, Gil, Dufayard, Dessimoz, & Gascuel, 2011). To obtain an estimate of divergence times between mtDNA haplotypes, we inferred a time-calibrated tree using BEAST v. 2.5.0. (Bouckaert et al., 2019). Based on results from previous work (Santos et al., 2014), we set a log-normal prior with mean 8.13 million years (MY) and standard deviation 1.2 MY (i.e. log(mean) = 2.12, log(s.d.) = 0.1) for the root age of *Phyllobates*. We used a Calibrated Yule tree prior, and set default priors for all other parameters, except for the clock rate mean and the Yule birth rate, which were set to *gamma*(0.01, 1000). We ran the MCMC sampler for 100 million iterations, sampling every 10,000, and generated a maximum clade credibility (MCC) tree using Tree Annotator (distributed with BEAST) after discarding the first 5% of trees as burnin. Mixing and stationarity of BEAST and MrBayes runs were evaluated visually and based on effective sample sizes (ESS) using Tracer v. 1.5 (Rambaut & Drummond, 2009). All mtDNA analyses were performed under partitioning schemes and molecular evolution models chosen with PartitionFinder2 (Lanfear, Frandsen, Wright, Senfeld, & Calcott, 2017).

### Transcriptome-enabled exon capture

Based on the results of mtDNA analyses we chose 63 samples (60 ingroup, 3 outgroup) from 17 localities (Table S1) representing the range of observed mtDNA variation among Colombian populations, and used them to perform transcriptome-enabled exon capture (Bi et al., 2012; Hodges et al., 2007). Briefly, we designed a set of DNA capture probes based on a transcriptome assembly and used them to enrich sequencing libraries for a subset of the genome.

#### Transcriptome sequencing

We generated a transcriptome assembly from liver, muscle, skin, and heart tissue of a single *P. bicolor* juvenile (NCBI BioSample SAMN15546883). RNA was extracted using Qiagen RNeasy spin columns, and pooled in equimolar ratios by tissue type to build a single cDNA library, which was sequenced on an Illumina HiSeq 2000. We filtered and trimmed reads using Trimmomatic v. 0.25 (Bolger, Lohse, & Usadel, 2014), and used Trinity (release 2013-02-25; Grabherr et al., 2011) to assemble them under default parameters, except for the minimum contig length, which was increased to 250bp. Finally we collapsed redundant contigs (e.g. alternative isoforoms) with CD-HIT-EST V.4.5.3 (Fu, Niu, Zhu, Wu, & Li, 2012).

#### Enrichment probe design

We annotated our transcriptome using BLASTX (Altschul, 1997) against *Xenopus tropicalis* proteins (JGI 4.2.72), and used Exonerate (Slater & Birney, 2005) to identify intron-exon boundaries in order to split transcripts into individual exons. We then chose a final set of exons to enrich in the following way: First we discarded those under 100bp, with GC content below 40% and above 70%, or which overlapped by more than 10bp based on Exonerate annotations. Next, we identified putatively repetitive elements and RNA-coding genes (e.g. rRNAs) in our transcriptome assembly with RepeatMasker v. 4.0 (Smith, Hubley, & Green, 2013) and BLASTn, respectively, and removed exons overlapping them. Finally, we blasted our exon set against itself with BLASTn under default parameters, and whenever two or more exons matched each other (e-value < 10^−10^), we retained only one of them. This resulted in 38,888 exons (7.57Mb) that passed filters, which were used to design 1,943,120 100bp probes that were printed on two Agilent SureSelect custom 1M-feature microarrays (3bp tiling).

#### DNA library preparation, target enrichment, and sequencing

We extracted DNA as described above, and used a Diagenode Bioruptor to shear each extraction to a ~100-500bp fragment distribution by performing 3-4 rounds of sonication (7min of 30s on/off cycles per round). DNA libraries were built following Meyer & Kircher (2010), except for bead cleanups, where we used a 1.6:1 ratio of beads to library (1.8:1 is recommended) to obtain a slightly larger final fragment size distribution. Finished libraries were combined in equimolar ratios into two 22.5 μg pools (one per array) for target enrichment. Array hybridization was performed largely following Hodges et al. (2009) with minor modifications: Each library pool was mixed with xGen Universal P5 and P7 blocking oligonucleotides and a mixture of chicken, human, and mouse COT-1 DNA. The two capture eluates were amplified separately by 18 cycles of PCR. To reduce the propagation of PCR-induced errors, each eluate was amplified in four parallel reactions. PCR products were pooled so that both captures were equally represented, and sequenced on Ilumina HiSeq 2500 and 4000 machines.

#### Bioinformatic pipeline

De-multiplexed read files were filtered by collapsing PCR-duplicate reads with SuperDeduper (Petersen, Streett, Gerritsen, Hunter, & Settles, 2015), trimming low quality bases and removing adapter contamination with Trimmonatic (Bolger et al., 2014) and Skewer (Jiang, Lei, Ding, & Zhu, 2014) under default parameters, except for the minimum read length, which was increased to 36 bp, and merging overlapping read pairs with FLASH (Magoč & Salzberg, 2011). To generate a reference for read mapping, we combined all cleaned reads from the ingroup species (i.e. *P. terribilis*, *aurotaenia*, and *bicolor*), and generated six *de novo* assemblies with different kmer sizes (k = 21, 31, 41, 51, 61, and 71) using ABySS (J. T. Simpson et al., 2009). We then merged the six assemblies using CD-HIT-EST and Cap3 (Huang & Madan, 1999). Finally, we identified contigs that matched our target exons using BLASTn, and retained only these for further analyses.

Reads from each sample were mapped to the reference using Bowtie2 v. 2.1.0 (Langmead & Salzberg, 2012), and outputs were sorted with Samtools v. 1.0 (Li et al., 2009), de-duplicated with Picard v.1.8.4 (http://broadinstitute.github.io/picard), and re-aligned around indels with GATK v. 3.3.0 (McKenna et al., 2010). We filtered our data in the following ways: First, we performed a reciprocal blast using the methods described above and removed any contigs with more than one match (e-value<10^−10^).

Second, we used ngsParalog (https://github.com/tplinderoth/ngsParalog) to identify contigs with variants stemming from read mismapping due to paralogy and/or incorrect assembly. This program uses allele frequencies to calculate a likelihood ratio for whether the reads covering a site are derived from more than one locus in the genome, while incorporating the uncertainty inherent in NGS genotyping. We calculated p-values for these likelihood ratios based on a 50:50 mixed χ^2^ distribution with one and zero degrees of freedom under the null, and removed any contigs with significantly paralogous sites after Bonferronni correction (α = 0.05). Third, we restricted all analyses to contigs covered by at least one read in at least 20 individuals, bases with quality above 30, and read pairs mapping uniquely to the same contig (i.e. proper pairs) with mapping quality above 20. Finally, we removed samples with less than 2.5 million sites covered by at least one read after filtering. This resulted in a dataset of 32,516 contigs (12.95 Mb) and 57 samples, which were used in all downstream analyses.

### Population Structure

To characterize genome-wide patterns of population differentiation we used our exon capture dataset to perform Principal Component Analysis (PCA) of genetic covariances calculated in PCangsd v.0.94 (Meisner & Albrechtsen, 2018), to estimate admixture proportions (k = 2-9) in ngsAdmix v.32 (Skotte, Korneliussen, & Albrechtsen, 2013), and to build a minimum-evolution tree in FastME v.2.1.5 (Lefort, Desper, & Gascuel, 2015) using genetic distances estimated with ngsDist (Vieira, Lassalle, Korneliussen, & Fumagalli, 2016). Nodal support for this tree was evaluated using 500 bootstrapped distance matrices produced in ngsDist by sampling blocks of 10 SNPs. These three analyses used genotype likelihoods (GL) as input, which were estimated in Angsd v.0.9.18 (Korneliussen, Albrechtsen, & Nielsen, 2014) at sites covered by at least one read in at least 50% of the samples without filtering for linkage disequilibrium. PCA and ngsAdmix analyses used one site per contig, randomly chosen among those with minor allele frequencies above 0.05 that passed the programs’ internal quality filters (5,634 sites). Genetic distance estimation for the ME tree was restricted to variable sites (i.e. SNP p-value < 0.05; 84,218 sites). PCA and ngsAdmix were run only on Colombian samples while the ME tree also included outgroups.

Finally, we reconstructed a population graph to evaluate historical splits and mixtures between sampling localities using Treemix (Pickrell & Pritchard, 2012). We called genotypes using the HaplotypeCaller and GenotypeGVCFs tools of GATK v.3.3.0 under default parameters, except for the heterozygozity prior, minimum base quality, and minimum variant-calling confidence, which were increased to 0.005, 30, and 20, respectively, to accommodate for the multi-species nature of our dataset. We then obtained allele counts for biallelic SNPs that were at least 1kb apart within each contig (usually resulting in a single SNP per contig, since most contigs were under 1kb), and with at least 50% genotyping (20,275 SNPs), using Plink v.1.90 (Purcell et al., 2007). In two cases, two nearby populations of the same color pattern (16.4 and 19.7 Km apart; Fig S1), which clustered closely in all other genetic structure analyses, were merged into single demes for allele count estimation due to small sample sizes. In addition, since we only had exon capture data for one *P. vittatus* individual, only *P. lugubris* was used as outgroup in this analysis. We ran Treemix v.1.13 assuming *m* = 0–6 migration edges, and chose the optimal number of migration edges by performing likelihood ratio tests in which we compared each value of *m* to the one immediately smaller. P-values were calculated based on a χ^2^ distribution with two degrees of freedom, since adding an extra edge adds two parameters (weight and direction of migration) to the model. This approach recovered m=2 as the most likely scenario (Table S2); results for m=0-6 are presented in Fig. S2.

### Phylogenetic relationships between lineages

To reconstruct the phylogenetic relationships between *Phyllobates* lineages, we inferred a species tree under the multispecies coalescent model, assuming independent sites, as implemented in SNAPP (Bryant, Bouckaert, Felsenstein, Rosenberg, & RoyChoudhury, 2012). SNAPP requires individuals to be assigned to operational taxonomic units (OTUs) *a priori.* Given our small sample sizes for some localities, as well as the evidence of gene flow between localities (see Results section), we took an *ad-hoc* approach and grouped our sampling localities into eight geographically and phenotypically coherent groups that showed evidence of being genetically distinct entities (see locality colors in Fig. 2B). Briefly, each OTU contained individuals that were geographically close, displayed the same color pattern, and showed evidence of genetic clustering in population structure analyses. We did not require OTUs to be fully reproductively isolated from each other. Further details on our OTU selection criteria can be found in the online supplement.

For computational efficiency, SNAPP was run on a reduced version of the Treemix dataset described above, restricted to SNPs genotyped for at least 75% of individuals and at least one member of each OTU (5,938 SNPs). We ran the MCMC sampler under default priors for 1,000,000 iterations, sampling every 250, and discarded the first 150,000 as burnin. Stationarity and mixing were evaluated in Tracer (Rambaut & Drummond, 2009) as detailed above, and the posterior tree distribution was summarized as a maximum clade credibility (MCC) tree in TreeAnnotator. To obtain estimates of divergence times between OTUs, we assumed a mutation rate of *μ* = 1e^−9^ mutations per site per year (Crawford, 2003; Sun et al., 2015), and a generation time of one year (*Phyllobates* frogs are sexually mature at ~10-18 months after hatching; Myers et al., 1978; R. Márquez *pers obs.*), and converted branch lengths to time units as *T* = (*τg*/*μ*), where *T* is the divergence time in years, *τ* the branch length in coalescent units, *g* the generation time, and *μ* the mutation rate (Bryant et al., 2012).

### Phylogenetic Comparative Analyses

In order to evaluate whether the central or leapfrogging populations exhibit a derived color pattern, we performed ancestral state reconstruction along the SNAPP MCC tree using maximum parsimony (Fitch, 1971) in the R package *phangorn* (Schliep, 2011). Aposematic coloration has been shown to co-evolve with several other traits, such as body size, toxicity, and diet specialization in dendrobatid frogs (Pough & Taigen, 1990; Santos & Cannatella, 2011; Summers & Clough, 2001). Understanding correlations between these traits within *Phyllobates* can shed light on how predation pressures have affected the geographic distribution of color patterns in this group. Therefore, we investigated the extent of correlated evolution between color pattern, body size, and toxicity. We used the snout-to-vent length (SVL) as a proxy for body size, and the average amount of batrachotoxin (BTX) in a frog’s skin as a proxy for toxicity. BTX is the most abundant and toxic alkaloid found in *Phyllobates* skins (Märki & Witkop, 1963; Myers et al., 1978). BTX levels were obtained from Table 2 of Daly, Myers, & Whittaker (1987), and SVL was measured from specimens in natural history collections (193 specimens; Table S3). We used mean SVL values for each lineage in analyses, and log-transformed BTX levels to attain normality of residuals. Correlations between traits were evaluated using phylogenetic generalized least squares regression (pGLS; Grafen, 1989; Martins & Hansen, 1997) with either Brownian motion (Felsenstein, 1985), Lambda (Pagel, 1999), or Ornstein–Uhlenbeck (Martins & Hansen, 1997) correlation structures. The best correlation structure was chosen by performing pGLS with the three correlation structures and comparing the fit of each model based on the AIC. Correlation structures were generated using the R package *ape* (Paradis, Claude, & Strimmer, 2004), and regressions were performed in the *nlme* package (Pinheiro, Bates, DebRoy, & Sarkar, 2017). In addition to the highest clade credibility tree, we also conducted tests of phylogenetic correlations on 1,000 randomly selected trees from the post-burnin SNAPP posterior distribution to account for phylogenetic uncertainty.

### Spatial population genetics

We took a spatial population genetics approach to investigate the extent of divergence and gene flow between populations in a spatially explicit way, aimed at understanding the nature and drivers of genetic variation across the landscape. First we generated a geo-genetic map of the Colombian *Phyllobates* populations using SpaceMix (Bradburd, Ralph, & Coop, 2016). This consists of a bidimensional plot where the distances between populations correspond to their expected geographic distances under stationary isolation by distance (IBD). Differences between geographic and geo-genetic locations therefore reflect historical rates of gene flow across the landscape. Populations that exchange more alleles than expected under stationary IBD are closer in geo-genetic than geographic space, and vice versa. For example, populations separated by a topographic barrier will be further apart in geo-genetic than geographic space. As input for SpaceMix we used allele counts generated as detailed above (see *Population Structure* section), for sites that were variable among Colombian individuals (8,093 sites). We then parameterized the full (“source_and target”) SpaceMix model with an MCMC run comprised of 10 initial exploratory chains (500,000 iterations each), followed by a 500,000,000 iteration “long” run, which was sampled every 10,000 iterations. We used default prior settings, and centered spatial (i.e. location) priors for each population at their sampling location. For the two demes composed of individuals from nearby localities we used the midpoint of the segment connecting both localities (Fig. S1).

SpaceMix accounts for the fact that a fraction of a population’s alleles may have been acquired through recent long-range migration from another region of the map by incorporating a long-range admixture proportion parameter, labelled *w*, which represents the probability that an allele in a given population migrated recently from a distant region. Since leapfrog distributions can, in principle, arise through introgressive hybridization between disjunct populations, we evaluated the support of our data for models with and without long-distance gene flow between populations (ie. all *w* parameters set to zero vs. *w* allowed to vary). We did so by estimating the Bayes Factor (Kass & Raftery, 1995) between both models using the Savage-Dickey density ratio (Dickey & Lientz, 1970), which approximates the Bayes Factor between nested models. Further details on this estimator and our implementation for SpaceMix models can be found in the online supplement.

Next, we used EEMS (Petkova, Novembre, & Stephens, 2015) to identify areas of the landscape where gene flow between populations is especially prevalent or reduced. Briefly, this algorithm estimates the rate at which genetic similarity decays with distance (i.e. the effective migration). Regions where this decay is quick or slow can be interpreted as barriers or corridors of migration, respectively. We estimated mean squared genetic differences between samples from genotype likelihoods in ATLAS (Link et al., 2017), and used them as input for EEMS. We set the number of demes to 500, and averaged across 10 independent 10,000,000-step MCMC runs logged every 1,000 steps (20% burnin). Since our genetic dissimilarity matrix was inferred from genotype likelihoods, specifying the number of SNPs used to compute the matrix (required by EEMS) was not straightforward. We used the number of sites with SNP p-value below 0.01, as calculated with Angsd (221,825 sites).

Finally, we assessed how attributes of the landscape influence genetic divergence between populations. Based on the results of EEMS and SpaceMix analyses, we evaluated the effect of three landscape features on genetic divergence: geographic distance, differences in elevation, and the presence of the San Juan River as a potential corridor of gene flow. To do so, we used the multiple matrix regression with randomization (MMRR) approach proposed by Wang (2013), which is an extension of multiple linear regression for distance matrices.

Our regression model consisted of genetic distance as a response variable and geographic distance, difference in elevation, and the effect of the San Juan river as a dispersal corridor as explanatory variables. As a proxy for genetic distance, we used the linearized genome-wide weighted FST (FST/[1-FST]; J. Reynolds, Weir, & Cockerham, 1983; Weir & Cockerham, 1984), estimated using Angsd based on 2D-site frequency spectra (SFS). To maximize the number of sites used to estimate each SFS, we included contigs with data for less than 20 individuals (but that passed all other filters) in this analysis. We estimated geodesic distances among populations based on GPS coordinates taken in the field using the pointDistance() function of the raster R package (Hijmans, 2017), and calculated elevation differences based on measurements taken in the field or extracted from Google Earth. To generate a proxy for the San Juan river as a dispersal corridor we built a resistance layer where every pixel overlapping the San Juan river had a value of 1 and all others had a resistance value *R,* which made movement between two pixels along the San Juan *R* times more likely than between two pixels outside the river. We then used this layer to calculate least cost distances between populations with the costDistance() function of the gdistance R package (Dijkstra, 1959; van Etten, 2017). Finally, we regressed the least cost distance against the geodesic distance, and saved the model residuals as a measure of the component of the resistance distance not explained by geographic distance. These residuals were used as an explanatory variable in our model. Since setting a biologically realistic value for *R* was not straightforward, we performed five separate MMRR analyses using least-cost distances estimated with *R* = 2, 10, 20, 50, and 100. The MMRR analysis was run using the script archived by Wang (2013; https://doi.org/10.5061/dryad.kt71r) with 10,000 permutations to estimate p-values.

## Results

### *Population structure among Colombian* Phyllobates

As expected from a multi-species dataset, we found multiple genetically structured clusters of individuals, which were largely concordant across analyses of the exon-enrichment and mtDNA datasets (Fig. 2, Fig. S2-S3). However, these clusters align much more closely with geography than either coloration or the current taxonomy: All populations of *P. terribilis* grouped with the southern populations of *P. aurotaenia*, while the northeastern populations of *P. aurotaenia* clustered closely with the northern populations of *P. bicolor*. The southern *P. bicolor* and the *P. aurotaenia* populations east and west of the Baudó mountains also formed independent clusters, but their relationship to other lineages was less clear. Finally, two mtDNA sequences from captive-bred *P. aurotaenia* of unknown origin (sequenced by Grant et al. [2006] and Santos et al. [2009]) were sister to those from the southern populations of *P. bicolor* in our genealogy (Fig. 2A). These results highlight the existence of several previously unrecognized (i.e. cryptic) lineages. Notably, they reveal the existence of three independent solid-yellow lineages, instead of two as previously thought, since the populations currently classified as *P. bicolor* clustered as two clearly separate and independent lineages. This points to an even greater discordance between coloration phenotypes and genetic similarity than previously thought.

### Phylogenetic relationships and divergence times

The inferred species tree was generally consistent with our genetic structure results, since tree topologies largely mirrored geography: Most OTUs were sister to close geographic neighbors, and higher level relationships followed a north-south axis (Fig. 2-3). In addition, the three yellow lineages were recovered each as sister to a different striped lineage. The topology of the SNAPP tree was largely concordant with those obtained in mtDNA and Treemix analyses. We only found inconsistencies in the placement of the *P. aurotaenia* populations from the eastern and western flanks of the Serranía del Baudó: Mitochondrial haplotypes from these two populations were part of a closely-related clade that also included all *P. aurotaenia* sequences from the Atrato river, and this clade was sister to another one containing sequences from the northern *P. bicolor* and the San Juan *P. aurotaenia* (Fig. 2A). Treemix also recovered the eastern and western Baudó populations as sister taxa, but they were sister to the rest of the Colombian populations (Fig. 2D and S2). Finally SNAPP recovered only the western Baudó *P. aurotaenia* as sister to all other Colombian populations, while the eastern Baudó *P. aurotaenia* was sister to the southern *P. bicolor* (Fig. 3). Treemix inferred a migration edge from the base of the clade containing the populations of *P. bicolor* and *P. aurotaenia* from the San Juan and Atrato drainages into the eastern Baudó *P. aurotaenia* (Fig. 2D). Since Treemix reconciles instances where a bifurcating tree model, such as the one used by SNAPP, does not fit the data well by incorporating migration edges between branches of the tree, this result suggests that these differences may be due to gene flow among populations.

**Fig. 3.**
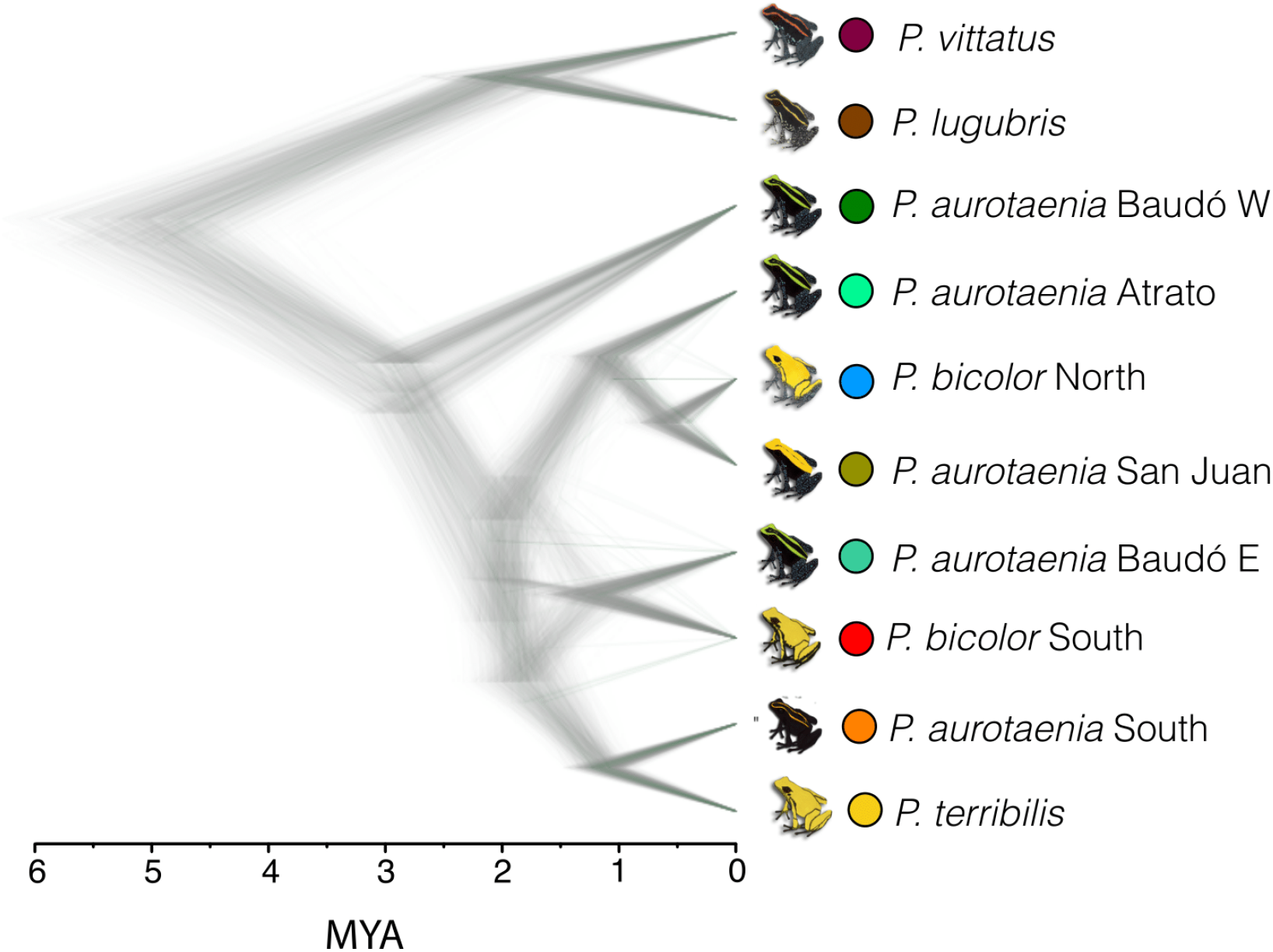
Phylogenetic relationships and divergence times among *Phyllobates* lineages inferred using SNAPP. Divergence times assume a mutation rate of 10^−9^ mutations per year and a generation time of one year. Each individual tree represents one sample from the SNAPP posterior distribution. Clades present in more posterior trees have higher posterior probabilities (i.e. higher nodal support). The color scheme is as in Fig. 2.

Divergence time estimation based on the SNAPP tree revealed a Plio-Pleistocene diversification of *Phyllobates*, and were generally concordant with previous estimates (Santos et al., 2009, 2014), indicating that our mutation rate and generation time assumptions are reasonable. The most recent common ancestor (MRCA) of *Phyllobates* was placed at 5.1 million years ago (MYA), with subsequent cladogenesis events from the late Pliocene to the Pleistocene (2.9-0.6 MYA; Fig. 3). These divergence times were slightly older but within the 95% HPD intervals of those estimated from mtDNA sequences (Fig. S4).

### Comparative analyses

Ancestral state reconstructions found the striped phenotype to be ancestral to solid-yellow (Fig. 4A). Phylogenetic regressions revealed a strong relationship between color pattern and size, with solid-yellow lineages being significantly larger than striped ones (Brownian Motion: β = 11.95, t = 9.92, df = 10, p = 9.03e-6; Fig. 4A), but a much weaker relationship between coloration and toxicity (Ornstein–Uhlenbeck: β = 1.19, t = 2.48, df = 5, p = 0.089; Fig. 4B). These results, suggest that at least two co-evolving traits (solid yellow coloration and larger size), possibly related to predator avoidance, are distributed in a leapfrog fashion in *Phyllobates*. Regressions performed over a set of posterior trees instead of the summary tree resulted in effect sizes and p-values centered around and qualitatively equivalent to those estimated using the summary tree, showing that the above conclusions are robust to the phylogenetic uncertainty present in our species tree reconstruction (Fig. S5).

**Fig. 4.**
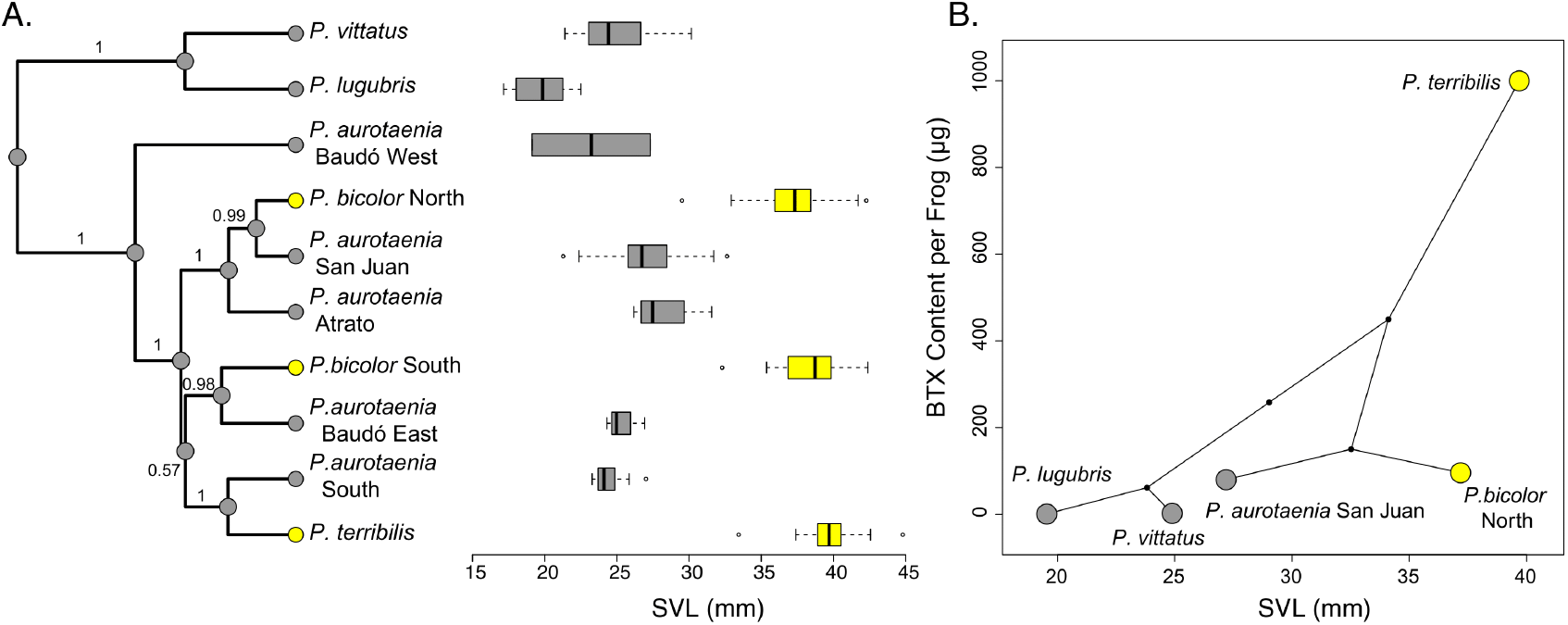
Evolutionary patterns of body size, color pattern, and toxicity in *Phyllobates*. Maximum clade credibility tree derived from the SNAPP posterior distribution and phylogenetic distribution of color pattern and snout-to-vent length (SVL) values among lineages. Numbers on internodes represent clade posterior probabilities. B) Phylogenetic biplot depicting the relationship between mean SVL and mean batrachotoxin concentration. Grey boxes/points represent striped lineages while yellow ones represent solid-yellow lineages.

### Spatial Population Genetics

The effective migration surface estimated by EEMS revealed a corridor of migration that matches the course of the San Juan river to a remarkable degree, considering that this method is completely agnostic to the topography of the landscape (Fig. 6A). This close match appears to lend strong support to to a high historical migration rate along the San Juan. However, we note that this result should be interpreted considering the sampling gap in the lower San Juan (See Fig. 5A). This corridor connects most of the sampled *P. aurotaenia* populations and the northern *P. bicolor*, and could explain the discordance between mtDNA and exon capture datasets in the phylogenetic placement of *P. aurotaenia* populations from the Eastern and Western Baudó mountains. Concordantly, SpaceMix estimated geo-genetic locations of populations along the San Juan corridor that were much closer to one another than their actual geographic positions: *P. aurotaenia* populations from the upper San Juan and Atrato drainages and the northern *P. bicolor* converged to very close locations in the upper/mid San Juan, overlapping considerably. The Baudó (east and west) and southern populations of *P. aurotaenia* were also shifted towards this area, but to a lesser extent (Fig. 5B).

**Fig. 5.**
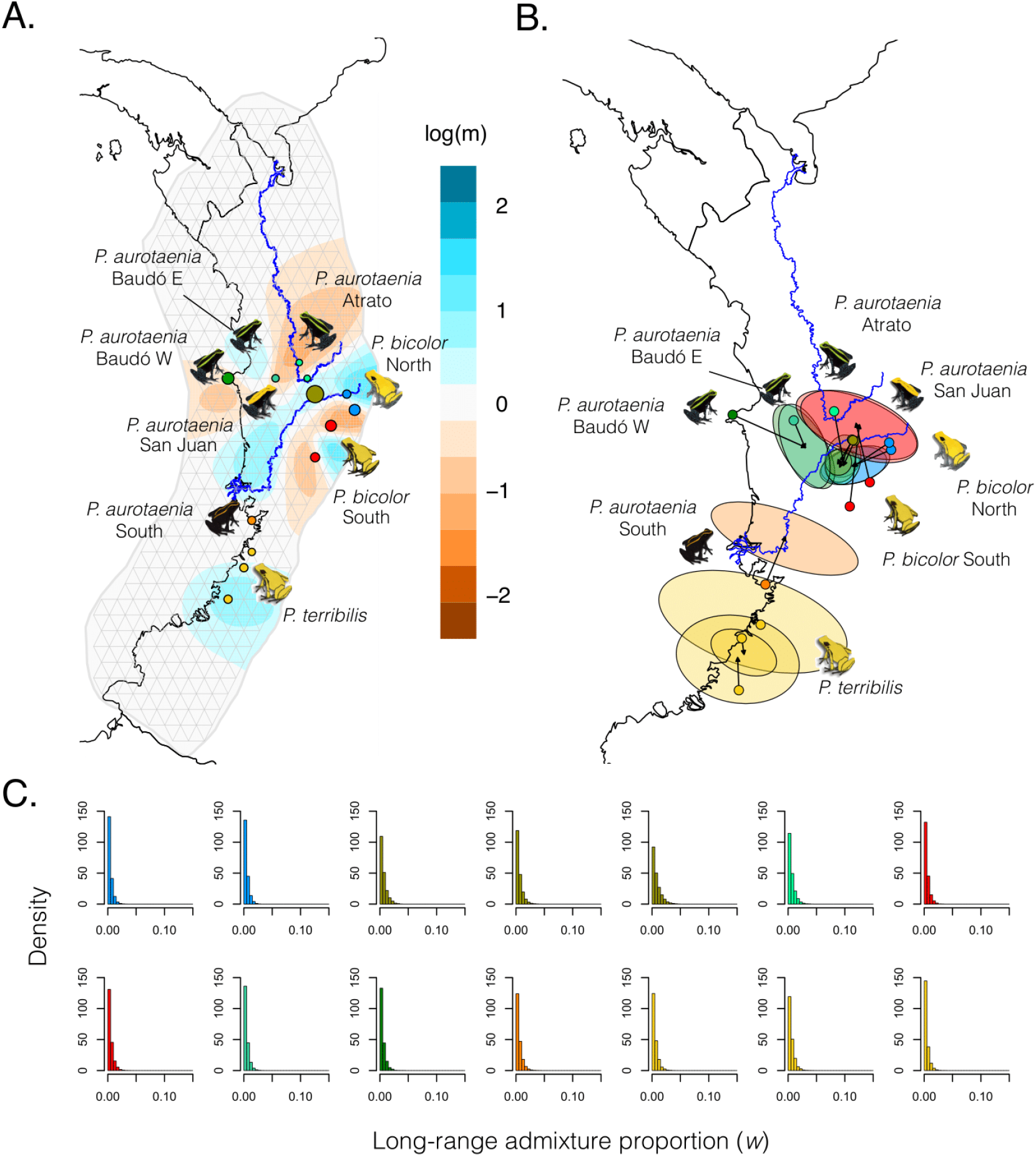
Gene flow among *Phyllobates* species across Western Colombia. A) Effective migration surface estimated using EEMS. Cyan and brown areas of the map are those where migration between demes is higher (cyan) or lower (brown) than expected under isolation by distance. Grey lines depict the population grid and habitat outline used by EEMS. B) Geo-genetic map inferred with SpaceMix. Ellipses represent the 95% Bayesian credible intervals around each population’s location on the geo-genetic map, and colored dots represent actual sampling locations. Arrows connect sampling and geogenetic locations. C) Density histograms of the posterior distributions of the long-range admixture proportion parameters (*w*) from the SpaceMix model for each population. Bars, points, and ellipses are colored by OTU as in Fig. 2. Histograms in Fig. 5C are ordered by OTU

In addition EEMS estimated very low levels of migration in the area enclosing the two southern *P. bicolor* populations, suggesting the existence of barriers to gene flow around these populations. Interestingly, the geo-genetic location of these populations was inferred north of its geographic location, past the mid San Juan cluster, and only slightly overlapping with other populations (Fig. 5B). The estimated long-distance admixture proportions were minimal for all populations (Fig. 5C), and the model with these proportions fixed at 0 was overwhelmingly supported over one where they were allowed to vary (Bayes factor = 1748).

In agreement with EEMS and SpaceMix results, the MMRR analysis found significant effects of geographic distance, elevation differences, and the San Juan as a migration corridor on genetic divergence between localities. Across the range of resistance values used, geographic distance was the strongest predictor. However, elevation differences and the San Juan as a barrier still had appreciable effects on genetic divergence (Table 1).

**Table 1.** Results from the multiple matrix regression with randomization (MMRR) analyses performed with multiple different resistance values for the San Juan river as a corridor (see Methods for details). P-values were estimated using 10,000 permutations. Coef. = coefficient; Diff. = difference; Dist. = distance; LC = least-cost.

| Predictor | Resistance = 100 |  |  | Resistance = 50 |  |  | Resistance = 20 |  |  | Resistance = 10 |  |  | Resistance = 1 |  |  |
| --- | --- | --- | --- | --- | --- | --- | --- | --- | --- | --- | --- | --- | --- | --- | --- |
|  | Coef. | t | p | Coef. | t | p | Coef. | t | p | Coef. | t | p | Coef. | t | p |
| Intercept | 0.0325 | 0.5286 | 1 | 0.0325 | 0.5298 | 1 | 0.0327 | 0.5343 | 1 | 0.0338 | 0.5427 | 1 | 0.0369 | 0.5953 | 1 |
| Geodesic Dist. | 0.6419 | 10.0547 | < 1e-4 | 0.6420 | 10.0618 | 0.0001 | 0.6423 | 10.0918 | < 1e-4 | 0.6428 | 10.1470 | < 1e-4 | 0.6483 | 10.0550 | < 1e-4 |
| San Juan LC Dist. | 0.2386 | 4.0094 | 0.0137 | 0.2392 | 4.0241 | 0.0138 | 0.2420 | 4.0890 | 0.0118 | 0.2471 | 4.2051 | 0.0118 | 0.2239 | 3.7483 | 0.0126 |
| Elevation Diff. | 0.2886 | 4.5994 | 0.0032 | 0.2877 | 4.5917 | 0.0037 | 0.2852 | 4.5697 | 0.0023 | 0.2802 | 4.5231 | 0.0023 | 0.2271 | 3.6051 | 0.0129 |
| | $r^2 = 0.62$, $F = 48.13$, $p < 1e-4$ | | | $r^2 = 0.62$, $F = 48.21$, $p < 1e-4$ | | | $r^2 = 0.63$, $F = 48.61$, $p < 1e-4$ | | | $r^2 = 0.63$, $F = 49.33$, $p < 1e-4$ | | | $r^2 = 0.62$, $F = 46.61$, $p < 1e-4$ | | |

## Discussion

Our main goal in this study was to understand the evolutionary and biogeographic processes that have shaped the leapfrog distribution of color pattern among *Phyllobates* populations, focusing on the roles of genetic divergence and gene flow. We found patterns of genetic structure and phylogenetic affinity between populations that closely match geography, evidence for gene flow between geographically close populations, especially along the San Juan river, and evidence against gene flow between distant populations.

These results provide strong evidence against the hypothesis that introgression of color pattern alleles between disjunct populations has played a role in generating the geographic distribution of this trait. Instead, they suggest an important role for short-range gene flow between neighboring populations. The high level of migration among the central striped populations along the San Juan river suggests that allele movement between these populations may have a homogenizing effect that maintains their phenotypic similarity. In addition, we find evidence for a barrier to gene flow that encloses the two sampled populations of the southern *P. bicolor* lineage, probably associated with differences in elevation, which could be helping maintain the phenotypic distinctiveness of this population. Conversely, the northern *P. bicolor* populations showed a strong signature of gene flow with their neighboring striped populations, suggesting that other forces, possibly selection, are maintaining the phenotypic differences between these populations in the face of recurrent gene flow. Nevertheless, to fully reject or accept these hypotheses, the history the alleles underlying color pattern differences must be taken into account (Hines et al., 2011).

Our phylogenetic reconstructions are consistent with a scenario in which the disjunct solid-yellow populations evolved their color patterns independently. However, the high levels of gene flow between geographically proximal populations and the close correspondence between phylogeny and geography lead us to suspect that, at least to an extent, the recovered phylogenetic relationships could be a product of prevalent gene flow between neighboring populations, and therefore may not reflect the history of color pattern evolution. A recent simulation study showed that even moderate levels of gene flow between geographic neighbors can confound phylogenetic inferences of convergent evolution (James et al., 2020). This scenario seems especially likely in the case of the northern populations of *P. bicolor*, given the signature of gene flow with their nearby striped populations (e.g. the San Juan and Atrato *P. aurotaenia*), but less so in the case of the southern *P. bicolor*, considering the strong barriers to gene flow inferred around the populations of this lineage. For *P. terribilis* we cannot favor either scenario, since we did not find strong evidence for or against gene flow with its sister *P. aurotaenia* populations.

It is therefore plausible that convergent evolution and common ancestry have both played a role in the origin of this leapfrog distribution. Teasing apart these two scenarios is challenging if gene flow between neighboring populations is prevalent, since high levels of genetic exchange between geographically close populations can erode existing differentiation between them, leading to patterns of genetic/phylogenetic affinity across the genome that mirror geography, regardless of their previous history. In the case of leapfrog distributions, this means that, in the face of persistent gene flow, peripheral populations will be closest to their phenotypically distinct neighbors, even if their phenotypic similarity stems from common ancestry. However, admixture between lineages is seldom uniform across the genome, since selection (see below) can restrict gene flow at certain genomic regions (J. R. Turner, Johnson, & Eanes, 1979; T. L. Turner, Hahn, & Nuzhdin, 2005; Wu, 2001). Such regions can therefore preserve historic signatures that have been erased by gene flow elsewhere in the genome. This is likely to be the case for loci underlying color pattern variation in *Phyllobates*, especially in cases such as the Northern *P. bicolor*, where phenotypic differences persist in spite of gene flow. Hence, the history of alleles at these loci should provide unique insights into the history of this phenotype. Future studies to identify such loci and understand their evolutionary history in relation to our current results will be instrumental to uncover the demographic processes leading to the current geographic and phylogenetic distribution of solid-yellow color pattern in *Phyllobates*, since they will allow for much more explicit tests of the hypotheses presented here.

Regardless of whether common ancestry or convergent evolution are at play in this system, it seems clear that differential selective pressures on the striped and solid-yellow populations have been involved in the origin and/or maintenance of the geographic distribution of color patterns. Independent evolution of similar phenotypes is many times promoted by similar changes in selective regimes (Darwin, 1859; Mayr, 1963; G. G. Simpson, 1953), and selection is required to maintain phenotypic differences between populations in the face of gene flow (Endler, 1977). Two of the three solid-yellow lineages (the northern and southern *P. bicolor*) occur at higher elevations (~600-1500 m.a.s.l) than the rest of the genus (~0-500 m.a.s.l). It is therefore possible that these mid-elevation habitats pose selective pressures (e.g. predator communities or light environments) different from those of lowland forests, which favor solid-yellow patterns over striped ones. The many known examples of variation in coloration across altitudinal gradients lend support to this idea (e.g. Köhler, Samietz, & Schielzeth, 2017; Rebelo & Siegfried, 1985; Reguera, Zamora-Camacho, & Moreno-Rueda, 2014; Richmond & Reeder, 2002; Rios & Álvarez-Castañeda, 2007). A similar situation could also be the case with *P. terribilis*, given its distribution at the southern edge of the genus’s range, where it may also experience different selective pressures from those faced by its closely related striped lineages.

The nature of color pattern variation in *Phyllobates* (i.e. solid vs striped) suggests that differential predation pressures may be important for the origin/maintenance of solid-yellow patterns. In aposematic species, advertisement signals with complex pattern elements, such as stripes, have been shown to serve a distance-dependent purpose, acting as conspicuous signals at short distances, while providing camouflage at long distances (Barnett et al., 2017; Barnett & Cuthill, 2014; Tullberg, Merilaita, & Wiklund, 2005). In contrast, bright, solid-colored signals remain conspicuous over a much wider range of distances. For example, a recent study focusing on the poison frog *Dendrobates tinctorius* found that striped patterns of this species are highly detectable at close range, but become camouflaged when observed from further away. Solid-yellow patterns, on the other hand, remained easily detectable over the whole range of distances tested (Barnett, Michalis, Scott-Samuel, & Cuthill, 2018). Therefore, it is likely that the striped and solid color patterns represent alternative aposematic strategies that are advantageous under different environments and/or predator communities.

The fact that we find a signature of correlated evolution between size and color pattern is compatible with this idea, since larger aposematic signals have been shown to be more detectable and memorable for predators (Forsman & Merilaita, 1999; Gamberale & Tullberg, 1996). Accordingly, size and conspicuousness are positively correlated among Dendrobatid poison frog species (Hagman & Forsman, 2003; Santos & Cannatella, 2011). However, we do not find a comparable pattern for toxicity, which has also been shown to co-vary with conspicuousness in poison frogs (Santos & Cannatella, 2011; Summers & Clough, 2001). This could be an artifact of low statistical power, since data are available only for one striped and two plain yellow Colombian lineages, but we cannot rule out the possibility that toxicity is indeed comparable between solid and striped populations. Furthermore, considering that aposematism relies on avoidance learning, it is possible that, despite similar levels of BTX, solid and striped populations differ in levels of palatability to predators. In any case, a scenario where solid and striped populations are similarly toxic and/or palatable is still compatible with predation pressures driving evolutionary convergence, since all species are considerably toxic (Daly et al., 1987; Myers et al., 1978). However, other explanations, such as geographic variation in mate preference (R. G. Reynolds & Fitzpatrick, 2007; Summers, Symula, Clough, & Cronin, 1999; Twomey, Vestergaard, & Summers, 2014; Yang, Richards-Zawacki, Devar, & Dugas, 2016), could also explain our results and cannot be ruled out.

It is worth noting, however, that the correlated evolution of body size and color pattern could also be due to ontogenetic integration (Olson & Miller, 1958). Tadpoles of all *Phyllobates* species are dark grey, and all of them develop a dorsolateral stripe shortly before metamorphosis, which remains unchanged until adulthood in striped lineages. Solid-yellow frogs, on the other hand, gradually lose dark pigmentation, until the solid adult pattern is attained a few months after metamorphosis (Myers et al., 1978). Therefore it is possible that, for example, the evolution of an extended growth period could generate changes in both body size and color pattern. If this is the case, then the concerted evolution of advertisement signal and body size would not necessarily be evidence of striped and solid patterns representing alternative predator avoidance strategies.

Our divergence time estimates indicate that the diversification of *Phyllobates* has followed the Plio-Pleistocene history of the Central American and the Chocó bioregions. The first cladogenesis event in our tree, which divides the Central American and Chocoan taxa was inferred between 4.5-5.9 MYA, which coincides with previously identified increases in faunal migration between Central and South America at ~6 MYA (Bacon et al., 2015; Santos et al., 2009). Further branching within South American lineages occurred later than 3 MYA, after both the Atrato (Duque-Caro, 1990b, 1990a) and Tumaco (Borrero et al., 2012) basins emerged above sea level to form the current Chocoan landscape. The Pleistocene was characterized by recurrent climatic and environmental fluctuations, which have been proposed as major drivers of neotropical rainforest biodiversity (Baker et al., 2020; Haffer, 1969; Hooghiemstra & Van Der Hammen, 1998; Vanzolini & Williams, 1970). Although the central Chocó has traditionally been regarded as a relatively stable Pleistocene forest refuge throughout the Quaternary (Gentry, 1982; Haffer, 1967; Hooghiemstra & Van Der Hammen, 1998), a notion supported by multiple palynological studies (Behling, Hooghiemstra, & Negret, 1998; Berrío, Behling, & Hooghiemstra, 2000; González, Urrego, & Martínez, 2006; Jaramillo & Bayona, 2000; Ramírez & Urrego, 2002; Urrego, Molina, Urrego, & Ramírez, 2006), there is some evidence of fluctuations in sea level, temperature, fluvial discharge, and, to a lesser extent, precipitation throughout the Quaternary in this region (González et al., 2006; Urrego et al., 2006). Despite being less dramatic than those experienced by other tropical forests (e.g. Amazonia), these fluctuations appear to have been related to changes in vegetation, especially the extent of mangrove forests (González et al., 2006). This may have promoted periodic retractions of *Phyllobates* populations towards the San Juan, perhaps resulting in increased rates of gene flow among them. Future work to understand how climatic fluctuations over the Quaternary have shaped the distribution of suitable habitat for *Phyllobates* frogs should shed further light on the biogeographic history of this genus in Northern South America.

Finally, our findings have broad implications for the systematics of *Phyllobates*. First and foremost, this study provides definitive evidence that the populations currently grouped under *P. aurotaenia* represent multiple independently-evolving lineages, some of which have probably been reproductively isolated for enough time to warrant recognition as separate species. Furthermore, we find that *P. bicolor* is comprised of two well-structured lineages that may have evolved similar phenotypes independently. In addition, we find highly variable levels of mtDNA divergence within *P. lugubris* (0-4% 16S, 0.2-8% COI, and 0-5.7% Cytb uncorrected p-distances), which could also be due to the existence of cryptic species.

It is, therefore, evident that a thorough revision of *Phyllobates* systematics is warranted. At this point, however, we refrain from modifying the group’s current taxonomy for several reasons. First, the complex interplay between divergence and gene flow that we have found in this system makes species delimitation based on genetic structure and coloration impractical. Therefore, integrating multiple lines of evidence (e.g. coloration, genetic variation, alkaloid profiles, bioacoustic data, larval and adult morphology) that allow us to draw stronger conclusions on the strength of reproductive barriers between lineages is needed to disentangle species limits. Second, although our study represents a substantial increase in geographic sampling, there are still considerable gaps, such as the lower San Juan drainage, or the mid-elevation forests south of the distribution of *P. bicolor,* that need to be considered. The fact that the two captive-bred *P. aurotaenia* included in mtDNA analyses are sister to the southern *P. bicolor* (Fig. 2A) suggests that we have not yet sampled the full diversity of P*hyllobates* lineages in Colombia. Finally the holotype of *P. aurotaenia* was collected in Condoto, Chocó, which is considerably distant from any of our sampling localities (Fig. S6), and the type locality of *P. bicolor* is unknown (Myers et al., 1978). This situation poses nomenclature issues, since, even if a robust species delimitation were available, naming these species would not be straightforward until the type specimens of *P. bicolor* and *P. aurotaenia* can be confidently assigned to one of them. Further work with increased sampling, including type specimens, and integrating multiple lines of evidence is therefore still needed to generate a taxonomy for *Phyllobates* that more accurately represents the genus’s evolutionary history.

## Concluding remarks

Leapfrog distributions constitute geographic replicates for the occurrence of a phenotype, and therefore provide important information about the origins of phenotypic diversity among closely related lineages. Here we show that, despite marked genetic structure and differentiation, there is considerable gene flow between phenotypically similar populations at the center of a poison-dart frog leapfrog distribution. This has probably been important for the origin and maintenance of the geographic distribution of color patterns in this group. Furthermore, we found instances of both reduced and increased levels of gene flow between neighboring populations with different phenotypes, suggesting that in some cases reduced gene exchange can contribute to the maintenance of phenotypic differences between populations in a leapfrog distribution, while in others these differences actually persist in the face of gene flow, probably due to local adaptation of different forms.

However, we are unable to answer a commonly addressed question about leapfrog distributions: whether phenotypic differences between populations stem from common ancestry or independent evolution. Even though our phylogenetic reconstructions unambiguously suggest the latter on their own, our finding of extensive gene flow among neighboring populations casts doubt on this conclusion. Several other studies on the history of leapfrog distributions have obtained similar phylogenies that align with geography instead of phenotypic similarity (Cadena et al., 2010; Garcia-Moreno & Fjeldså, 1999; Norman et al., 2002; Quek et al., 2010; Toon, Austin, Dolman, Pedler, & Joseph, 2012), leading to the view that leapfrog distributions are often due to independent evolution. Our results, therefore, add to the notion that processes such as pervasive gene flow (Hines et al., 2011; James et al., 2020) or incomplete lineage sorting (Avise & Robinson, 2008) can decouple the history of phenotypic change at a given trait from genome-wide patterns of divergence, possibly leading to erroneous inferences of convergent evolution (Hahn & Nakhleh, 2015).

## Supporting information

Supplementary tables (one table per sheet)

## Data accessibility

mtDNA sequences were uploaded to GenBank under accessions MT742690-MT742754, MT749179-MT749246, and MT808222-MT808283. Raw Illumina reads were uploaded to the NCBI SRA under BioProject ID PRJNA645960. The assemblies, bam and vcf files, and body size data, as well as the code used for analyses are available at https://doi.org/10.5061/dryad.8d4r3vd or as supplementary material.

## Acknowledgements

We thank Pablo Palacios-Rodríguez, José Alfredo Hernández, Carolina Esquivel, Diana Galindo, Mabel González, and Fernando Vargas-Salinas for assistance in the field, Lydia Smith, Valeria Ramírez-Castañeda, Alvaro Hernández, and Ke Bi for help with molecular and bioinformatic procedures, Andrea Paz for advice on MMRR analyses, and Alan Resetar (FMNH), Andrew Crawford and Alberto Farfán (ANDES), Rayna Bell and Addison Wynn (USNM), Andrés Acosta and Carlos Montaña (IAvH), John Taylor Rengifo (UTCh), Greg Schneider (UMMZ), and David Kizirian (AMNH) for facilitating access to preserved specimens. Comments and suggestions from Trevor Price, John Novembre, Valentina Gómez-Bahamon, John Bates, Daniel Matute, Catalina Gonzalez, the Bates/Hackett lab, the Kronforst lab, and four anonymous reviewers greatly improved this paper. We sincerely thank Lina M. Arenas for allowing us to use her beautiful frog illustrations for our figures. This work was funded by a Basic Sciences Grant from the Vice Chancellor of Research at Universidad de los Andes, a Colombia Biodiversa Scholarship from the Alejandro Angel Escobar foundation, a Pew Biomedical Scholarship, Neubauer Family funds and a Steiner Award from the University of Chicago, and NSF grant DEB-1655336. RM was partially supported by a Fellowship for Young Researchers and Innovators (Otto de Greiff) from COLCIENCIAS. Computations were performed on the University of Chicago’s Gardner HPC cluster, funded by NIH grant TR000430. Tissue collections were authorized by permits No. 2194 and 1380 from the Colombian Ministry of Environment and Authority for Environmental Licenses (ANLA).

## Author contributions

R.M., A.A., and M.R.K. conceived the project, R.M., T.P.L, R.N., M.R.K, and A.A. designed the research, R.M., A.A., R.N. and M.R.K. acquired funding, A.A., D.M-V., and R.M. collected samples, R.M. and T.P.L. generated the data, and R.M. analyzed the data and wrote the paper with input from M.K. and edits from all authors.

## Supplementary information

**Supplementary note.** Criteria used to select Operational Taxonomic Units (OTUs) for phylogenetic inference with SNAPP.

**Table S1.** Information and accession numbers for samples used in this study. Locality data is limited to prevent illegal traffic. Further details are available from the authors upon request.

**Table S2.** Results of likelihood ratio tests performed on Treemix analyses run with m = 0-6 migration edges.

**Table S3.** Information and snout-to-vent lengths of museum specimens measured for comparative analyses.

**Figure S1.** Localities joined into a single deme for locality-level analyses.

**Figure S2.** Results of Treemix analyses run with m=0-6 migration edges.

**Figure S3.** Minimum-evolution tree based on genetic distances.

**Figure S4.** Mitochondrial DNA time tree inferred using BEAST 2.

**Figure S5.** Results of phylogenetic comparative analyses run on 1000 randomly-drawn trees from the SNAPP posterior tree distribution.

**Figure S6.** Map of the type locality of *P. aurotaenia*.

**Figure S7 and associated text.** Details on our implementation of the Savage-Dickey ratio to estimate Bayes Factors between nested SpaceMix models.

