## Supplementary figures and text for "Divergence, gene flow and the origin of leapfrog geographic distributions: The history of color pattern variation in *Phyllobates* poison-dart frogs"

### Supplementary Material

### Supplementary Note:
#### Criteria used to select Operational Taxonomic Units for SNAPP

Being a species tree inference algorith, SNAPP requires all gentoyped individuals to be assigned to an operational taxonomic unit (OTU), and infers the relationships and divergence times between these OTUs. Due to our low and non-uniform sample sizes across populations, the presence of some potentially important sampling gaps, and the strong evidence for gene flow across the landscape, we took an ad-hoc approach to OTU selection. We defined OTUs as geographically, genetically, and phenotypically cohesive groups that appeared to be independently evolving units.

More specifically, we assigned a group geographically close individuals to an OTU if they met at least three of the following criteria:

1. All individuals were either striped or solid-colored.
2. All individuals/localities were monophyletic in mtDNA, Treemix (Tmx), and minimum-evolution (ME) tree analyses.
3. Alternatively, all individuals/localities were paraphyletic with respect to members of at most one other OTU in mtDNA, Treemix, and minimum-evolution tree analyses.
4. All individuals formed a discrete cluster on the first three axes of the PCA, which did not overlap with individuals from any other group.
5. All individuals formed a distinct, unique block in ngsAdmix analyses for at least two values of K.

Below we detail criteria fulfilled by each of our eight OTUs:

| OTU | Color code | Criteria fulfilled |
|---|---|---|
| *P. aurotaenia* Baudó West | (dark green) | 1, 2a (mtDNA, ME), 3, 4 |
| *P. aurotaenia* Baudó East | (teal green) | 1, 2a (mtDNA), 2b (ME), 3 |
| *P. aurotaenia* Atrato | (bright green) | 1, 2a (mtDNA, ME), 3 |
| *P. bicolor* North | (blue) | 1, 2a (Tmx, ME), 2b (mtDNA), 4 |
| *P. aurotaenia* San Juan | (olive) | 1, 2b (Tmx, mtDNA, ME), 4 |
| *P. bicolor* South | (red) | 1, 2a (Tmx, mtDNA, ME), 3, 4 |
| *P. aurotaenia* South | (orange) | 1, 2a (mtDNA), 2b (ME), 3 |
| *P. terribilis* | (yellow) | 1, 2a (Tmx, mtDNA), 2b (ME), 3 |

Finally, we note that, since we obtained largely congruent phylogenetic results between our SNAPP analysis using these OTUs and other analyses that used individuals or sampling localities as the unit of analysis (ie. the mtDNA, ME, and Treemix trees), we are confident that our SNAPP results, and those derived from them (ie. phylogenetic comparative analyses) are biologically sound.

### Supplementary Figures

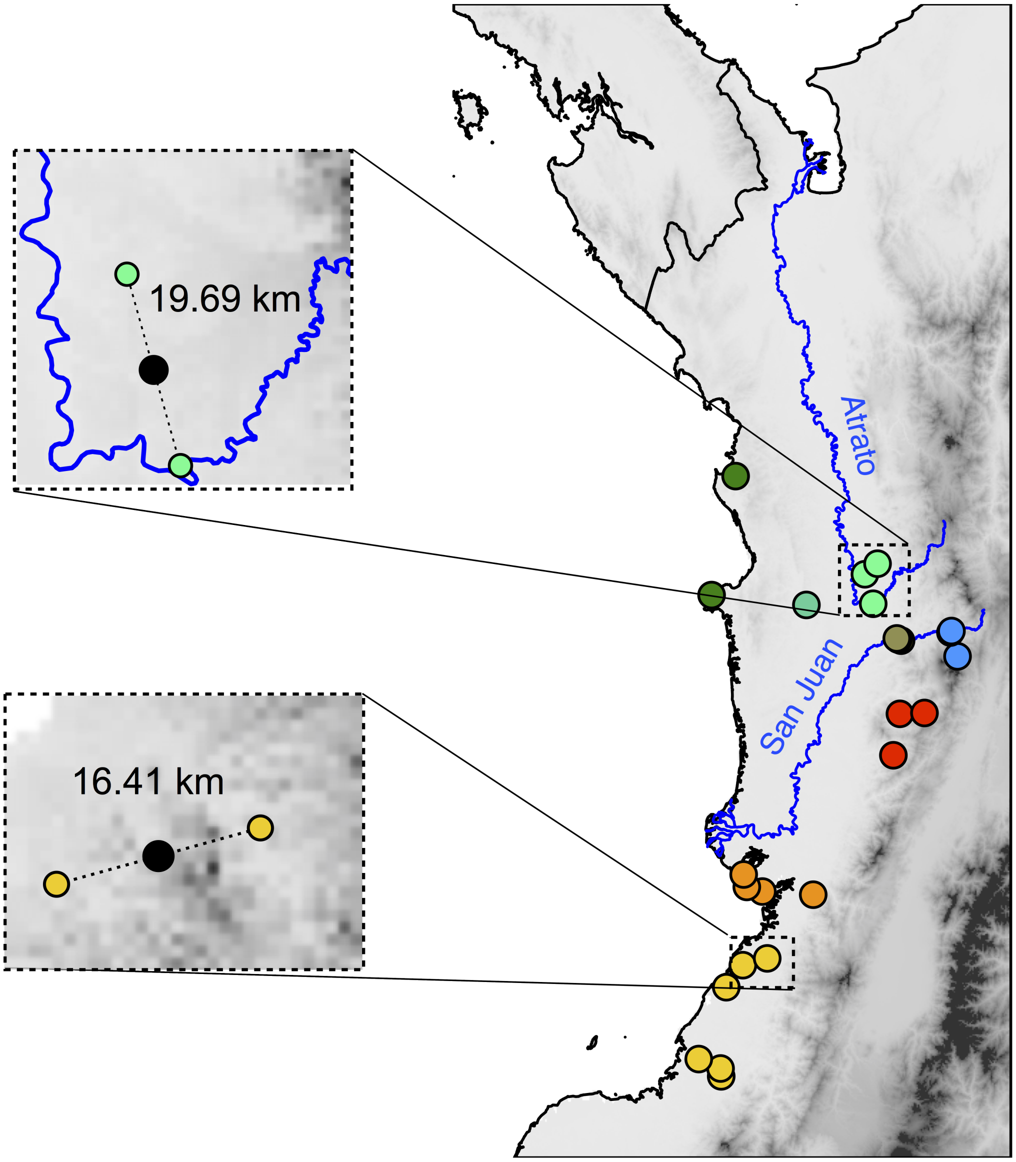


Figure S1: Localities joined into a single deme for Treemix, SpaceMix, and MMRR analyses analyses.

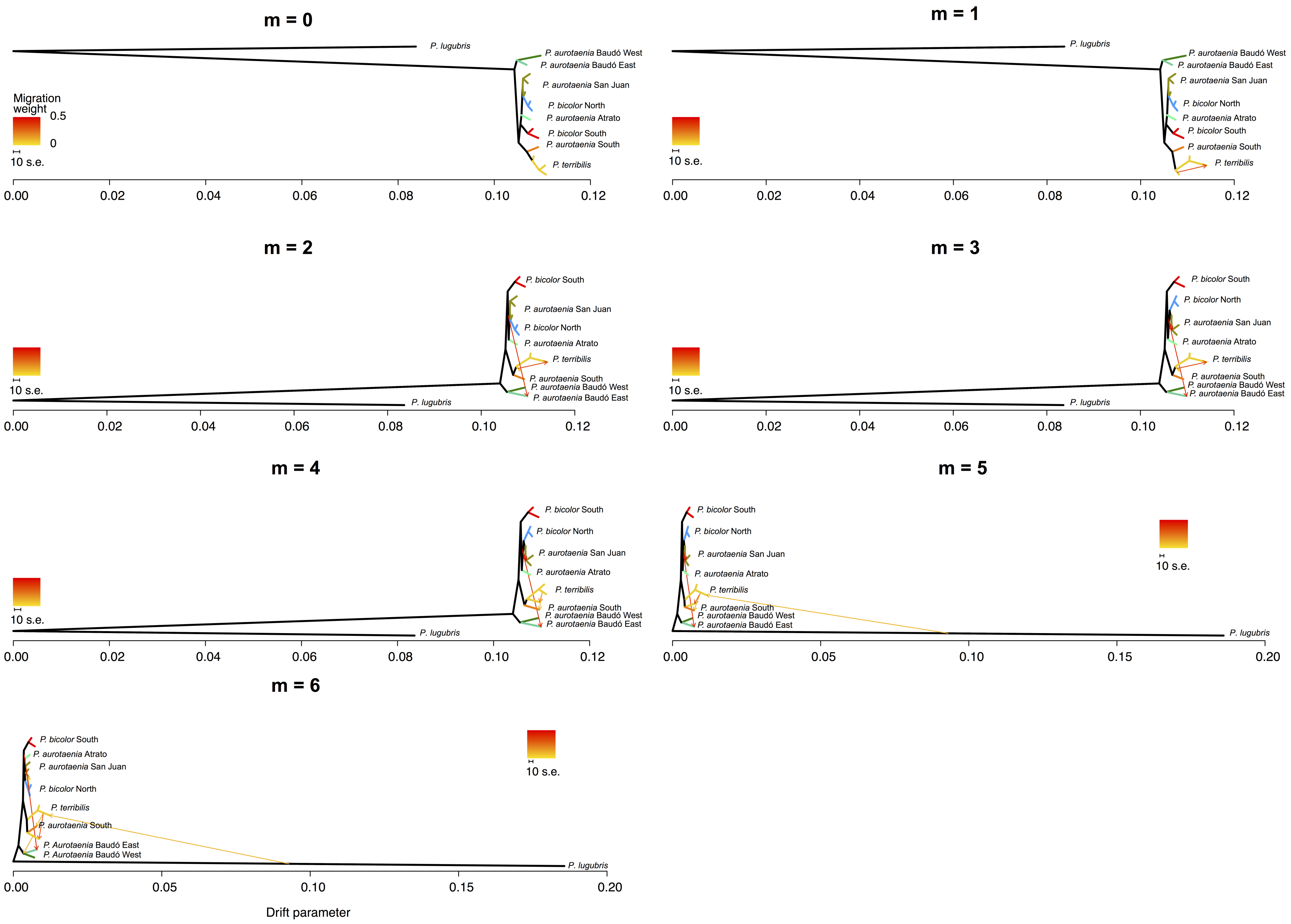


Figure S2: Results of Treemix analyses run with $m = 0 - 6$ migration edges.

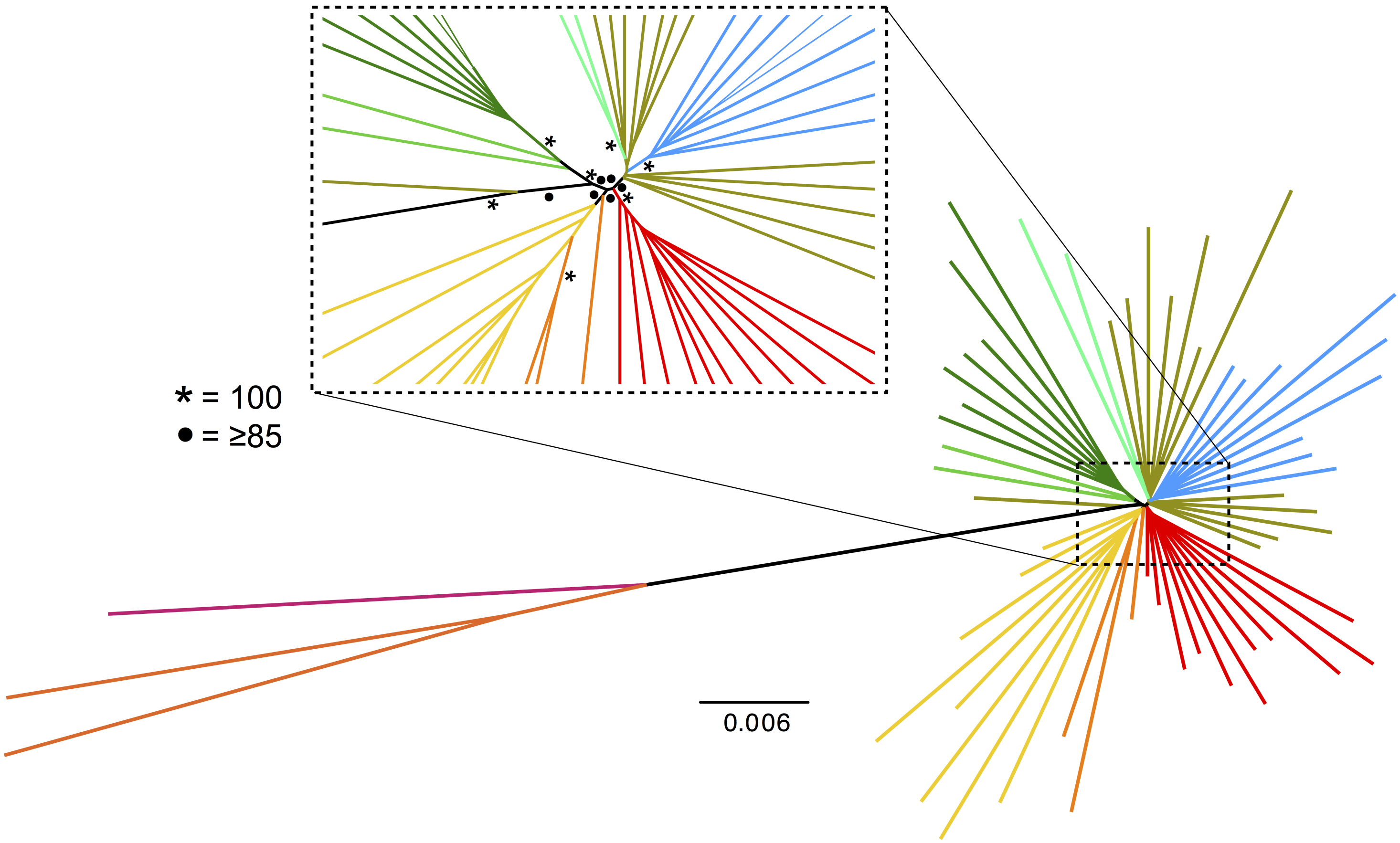


Figure S3: Minimum-evolution tree based on genetic distances. Symbols on internodes correspond to bootstrap support values.

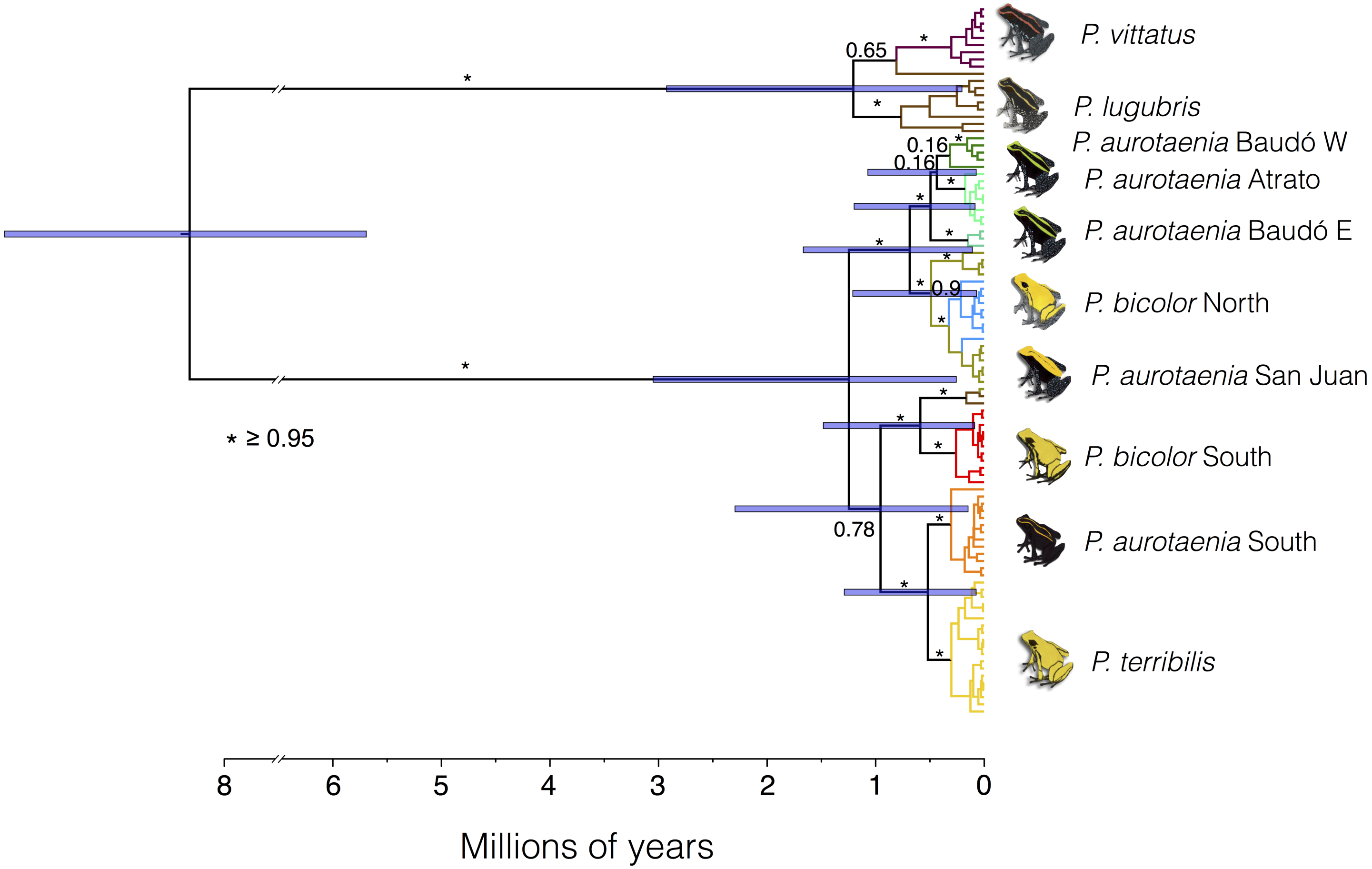


Figure S4: Mitochondrial DNA time tree inferred using BEAST 2. Symbols and numbers on internodes represent posterior probabilities in support of the following node, and purbple bars represent 95% highest posterior density intervals for node dates.

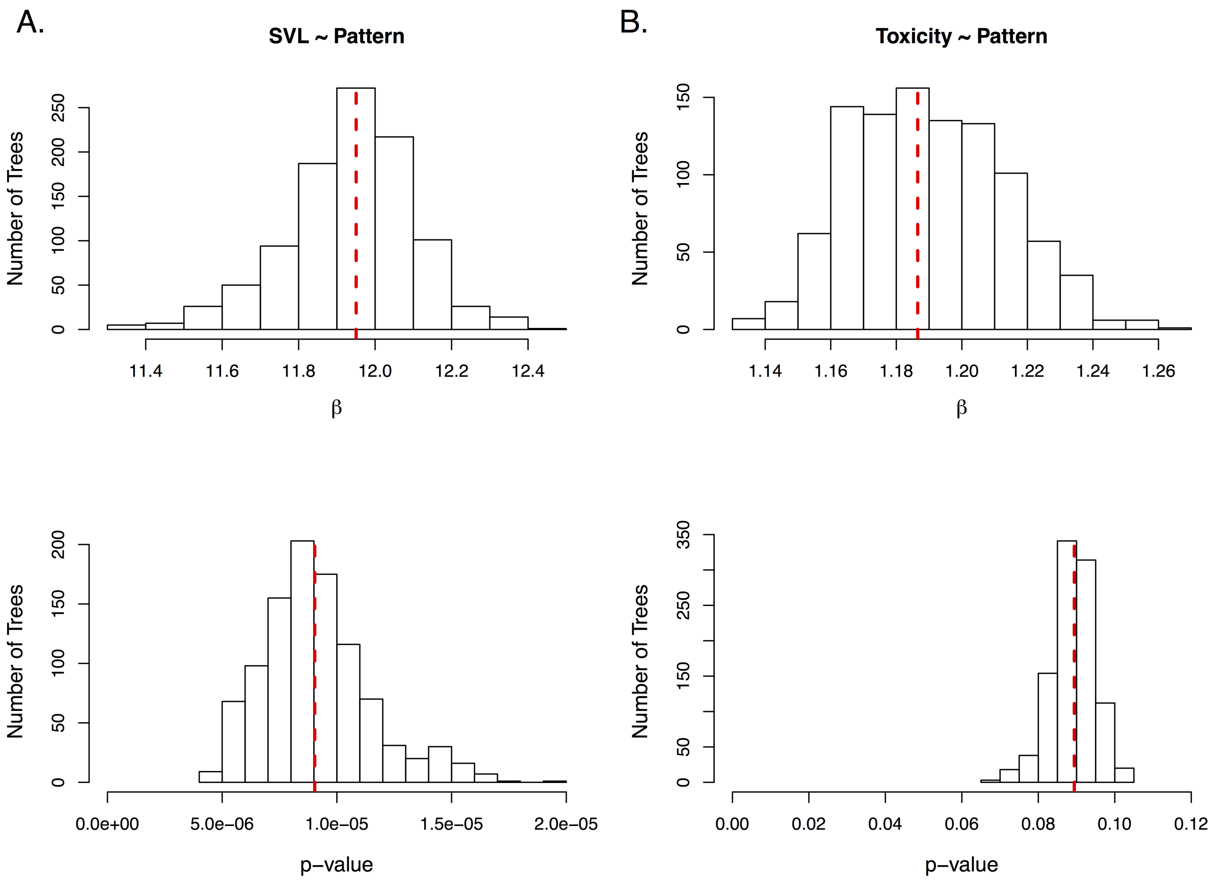


Figure S5: Results of pGLS regressions run over the SNAPP posterior tree distribution. A) regression of Pattern and SVL (i.e. body size) and B) BTX content (i.e. toxicity) and pattern. Histograms show the distribution of effect sizes ($\beta$) and p-values obtained by pGLS regressions run on 1000 randomly selected post-burnin trees. Vertical lines show the estimates obtained using the highest clade credibility tree, which are reported in the main text.

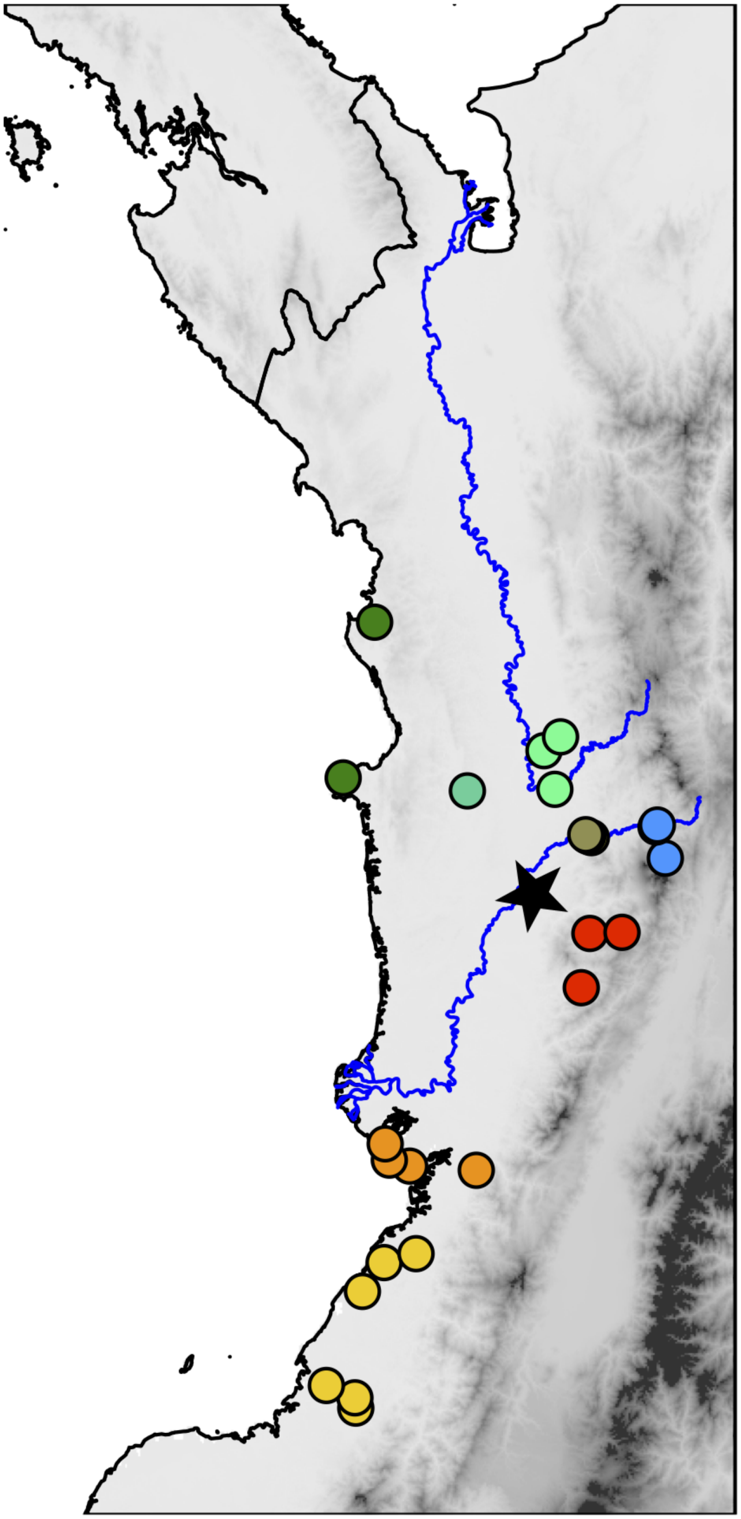

Figure S6: Map of the type locality of *P. aurotaenia* (star). Other localities are the same as in Fig. 2B of the main text.

### SpaceMix model comparison using the Savage-Dickey ratio to estimate Bayes Factors

*Bayes Factors*

A commonly used metric for model comparison in a Bayesian framework is the Bayes Factor (Kass & Raftery, 1995; Jeffreys, 1938), which is the ratio of the marginal likelihoods of two competing models. From Bayes's theorem, the ratio of the posterior probabilities of such models, $M_0$ and $M_1$, after observing data $D$ (i.e. the posterior odds) can be expressed as

$$\frac{P(M_0|D)}{P(M_1|D)} = \frac{P(M_0)}{P(M_1)}\frac{P(D|M_0)}{P(D|M_1)} \tag{1}$$

This expression shows how the Bayes Factor $[P(D|M_0)/P(D|M_1)]$ drives the change from prior odds $[P(M_0)/P(M_1)]$ to posterior odds $[P(M_0|D)/P(M_1|D)]$, providing an intuitive metric of how well each model explains the data.

The marginal likelihood of a model corresponds to the likelihood for that model, weighed by its priors, and integrated over all the parameter space:

$$P(D|M) = \int P(\theta|M)P(D|\theta, M)d\theta \tag{2}$$

Except for some simple models where analytical solutions are possible, estimating marginal likelihoods can easily become a large computational endeavor requiring complex MCMC sampling, especially for complex models with many parameters [see Marin & Robert (2009) and Oaks *et al* (2019) for recent reviews of available approaches].

*The Savage-Dickey ratio*

An interesting exception to the problem described above are comparisons between nested models, where one of the models being compared can be obtained by setting a subset of parameters of the other model to fixed values. More formally, if model $M_1$ has parameters $\theta = (\psi, \phi)$, where $\psi$ and $\phi$ are parameter vectors, model $M_0$ is considered to be nested within $M_1$ if $M_0$ has parameters $(\psi, \phi = \phi_0)$, where $\phi_0$ is a vector of fixed values, and if the prior and marginal likelihood of $M_1$ are equal to those of $M_0$ when $\phi$ is equal (or tends to) $\phi_0$. That is: $lim_{\phi\to\phi_0}P(\psi|\phi, M_1) = P(\psi|M_0)$ and $P(D|\langle\psi, \phi_0\rangle, M_1) = P(D|\psi, M_0)$.

Consider the marginal likelihood of $M_0$

$$P(D|M_0) = \int P(\psi|M_0)P(D|\psi, M_0)d\psi \tag{3}$$

If the conditions described above hold, this can be re-written as

$$P(D|M_0) = \int P(\psi|\phi_0, M_1) P(D| \langle \psi, \phi_0 \rangle, M_1) d\psi = P(D|M_1, \phi = \phi_0) \tag{4}$$

Applying Bayes's theorem

$$P(D|M_0) = \frac{P(\phi = \phi_0|D, M_1) P(D|M_1)}{P(\phi = \phi_0|M_1)} \tag{5}$$

From this we can obtain the Bayes factor in favor of $M_0$

$$\frac{P(D|M_0)}{P(D|M_1)} = \frac{P(\phi = \phi_0|D, M_1)}{P(\phi = \phi_0|M_1)} \tag{6}$$

This constitutes the ratio of the posterior and prior distributions of $\phi$ under $M_1$ evaluated at $\phi_0$, also known as the Savage-Dickey density ratio (Dickey & Lientz, 1970). Intuitively this ratio can be interpreted as the change from prior to posterior probability of $\phi = \phi_0$ after allowing parameters $\phi$ to vary.

*Application to SpaceMix models*

One of the main advantages of SpaceMix is that it explicitly incorporates admixture between populations in an isolation by distance context (Bradburd *et al*, 2016). This, among other things, allows users to test the extent to which long distance admixture explains the genetic covariance observed among populations by comparing models where populations draw admixture with those where the admixture proportions are fixed to 0. Since the second kind of model is nested within the first, the Savage-Diceky ratio can readily be used for this comparison as detailed below.

The parameters of interest for this comparison are the admixture proportions for each population $w = (w_1, w_2, \ldots w_k)$ which represent the probability $(0 < w_k < 0.5)$ that an allele sampled from population $k$ is of admixed origin (i.e. it originated at a geogenetic location different from that of $k$). The Bayes factor in favor of a model with fixed admixture proportions $(M_0)$ versus a model where admixture proportions are free to vary $(M_1)$ can be calculated using the Savage-Dickey ratio as

$$\frac{P(D|M_0)}{P(D|M_1)} = \frac{P(w = z|D, M_1)}{P(w = z|M_1)} \tag{7}$$

Where $w = (w_1, w_2, \ldots w_k)$ is the vector of admixture proportions for $k$ populations and $z = (z_1, z_2, \ldots z_k)$ the set of fixed admixture proportions defined under $M_0$. If one is comparing admixture versus no admixture then $z$ is composed entirely of zeros, but this approach can be used with any set of fixed values.

$P(w = z|M_1)$ represents the prior probability of observing $z$ under $M_1$. SpaceMix uses a prior distribution of $w_k$ where $w_k \sim Beta(\alpha = 1, \beta = 100)$ (Bradburd *et al*, 2016). Therefore, under the assumption that admixture proportions are independent from one another, this probability is simply the product of the $Beta(1, 100)$ distribution evaluated at each value of $2z$

$$P(w = z|M_1) = \prod_{i=1}^{k} Beta(1, 100)\Big|_{2z_i} \tag{8}$$

Following the same logic, the posterior probability of observing $z$ given $M_1$ and data $D$ is

$$P(w = z|D, M_1) = \prod_{i=1}^{k} P(w_i = z_i|D, M_1)\Big|_{z_i} \tag{9}$$

Since there is no analytical solution for $P(w_i = z_i|D, M_1)$ it must approximated from the posterior distribution of $w_i$ obtained through MCMC sampling. Below we illustrate how this approach was used to evaluate the extent of long-distance admixture in *Phyllobates* frogs, including the R code used.

Our goal was to evaluate the extent to which events of long-distance admixture have influenced the genetic covariance among 14 populations of *Phyllobates* frogs distributed across the biogeographic Chocó of Western Colombia (see main text for further details). In statistical terms, we aimed to estimate the Savage-Dickey ration between the full SpaceMix model, which allows for populations to draw long distance admixture ($M_1$), and one where all 14 admixture proportions ($z$ in Eq. 7) were fixed at 0 ($M_0$).

The denominator of the Savage-Dickey ratio is straightforward to calculate using equation 8:

```
z <- rep(0,14)
prior <- prod(dbeta(2*z, shape1=1, shape2=100))
prior
#[1] 1e+28
```

To estimate the numerator we parameterized the full (i.e."source_and_target") SpaceMix model to obtain well-sampled estimates of the posterior distributions of the admixture proportions for our 14 populations (i.e. $w = (w_1, w_2, \dots w_{14})$). We then approximated the posterior density of each $w$ parameter at 0 by fitting a logspline density function (Stone *et al*, 1997) and evaluating it at $w_k = 0$. Finally, we applied Eq. 9 to obtain the numerator and calculate the Savage-Dickey ratio.

```
library(SpaceMix)
library(polspline)

mcmc <- load("Spacemix_Counts_Full_space_MCMC_output1.Robj")

w_eval=numeric(14)

for(i in 1:14){
        w <- admix.proportions[i,]
        spline <- logspline(w, lbound=0) # Fit logspline density
        w_at0 <- dlogspline(0,spline) # Evaluate at w=0
        w_eval[i] <- w_at0
        }

posterior <- prod(w_eval)

posterior
#[1] 1.748146e+31
```

Briefly, the above code loops through the sets of posterior samples for $w_1 - w_{14}$, fits a logspline density and evaluates it at zero, generating a set set of posterior probability estimates whose product constitutes an approximation of $P(w = z|D, M_1)$.

Finally, we can use Eq. 7 to get the Bayes factor in support of $M_0$

$$\frac{P(D|M_0)}{P(D|M_1)} = \frac{P(w = z|D, M_1)}{P(w = z|M_1)} = \frac{1.748 \times 10^{31}}{1 \times 10^{28}} = 1748.16 \quad (10)$$

The above indicates that $M_0$ is overwhelmingly favored by the data over $M_1$, indicating strong support in for a scenario where gene flow between distant populations is very low or non existent.

A complementary way to visualize this result is plotting the prior and posterior distributions of all $w$ parameters (Fig. S7). This allows us to see that in all but one case the posterior density of $w$ at $w = 0$ is much higher than the prior, indicating that observing the data greatly increases support in favor of a scenario with no long-distance admixture. The only exception is $w_1$, there the posterior density at 0 is about one half as high as the prior. However, the posterior density very quickly increases and surpasses the prior at $w_1 = 1.86 \times 10^{-7}$, and has 95% of its mass below 0.012, so, although it doesn't provide support in favor of $w = 0$, the estimate for $w_1$ is very low and compatible with a scenario of minimal long-distance admixture.

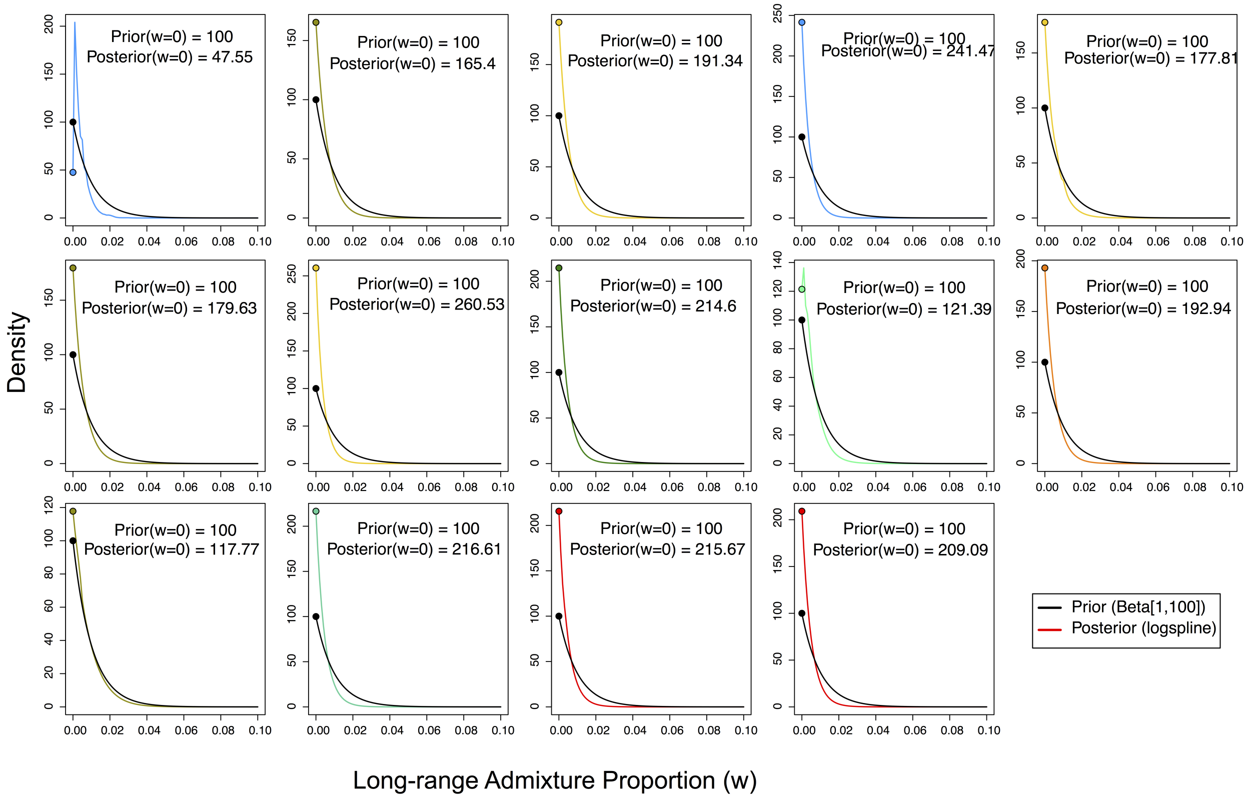


Figure S7: Prior and posterior (logspline) distributions of the admixture proportions for the 14 populations of *Phyllobates* analyzed. Prior distributions are colored black and posterior distributions as lines colored by OTU in accordance to Fig. 2 of the main text. Dots represent the point estimate of each distribution at $w_k = 0$.
